# whittle: annotation-preserving trimming and filtering of long reads

**DOI:** 10.64898/2026.11.27.769621

**Authors:** Erdi Kılıç, Burçak Otlu

## Abstract

**Summary:** Long reads from Oxford Nanopore and PacBio instruments can carry per-base annotations in unaligned BAM or FASTQ headers. These include base-modification calls, PacBio polymerase kinetics and Nanopore move tables that link bases to raw signal. Trimming and splitting remove bases or change their positions, so annotations that are not updated can refer to removed bases or the wrong positions. We present whittle, which removes adapters, barcodes and primers, splits chimeric reads, filters reads by quality and length, and rewrites the supported annotations of each retained segment. With supplied sequences, its mean base-level trimming F1 on simulated reads was 95.6% (ONT) and 95.9% (PacBio). In the tested reads, its move tables, retained modification calls and PacBio kinetic arrays matched values calculated independently from the untrimmed input. At eight threads, whittle cropped and filtered untagged gzip FASTQ 1.2 to 1.4 times as fast as chopper and 5.1 to 5.8 times as fast as fastplong.

**Availability and implementation:** whittle 1.0.0 is written in Rust and released under the Apache-2.0 license at https://github.com/erdikilic/whittle, with binaries for Linux and macOS and packages on Bioconda and crates.io. whittle 1.0.0 is archived at https://doi.org/10.5281/zenodo.24051287, and the benchmark data and scripts at https://doi.org/10.5281/zenodo.24051464.

**Supplementary information:** Supplementary data are available at *Bioinformatics* online.

Graphical abstract.
Reads from ONT and PacBio instruments enter whittle as uBAM or FASTQ after basecalling or consensus calling. Reads 1 and 3 are trimmed and kept, read 2 is removed as too short and read 4 for low quality, and read 5 is split at an internal adapter into reads 5a and 5b. Modification calls and per-base arrays stay with their bases. Instruments are not to scale.

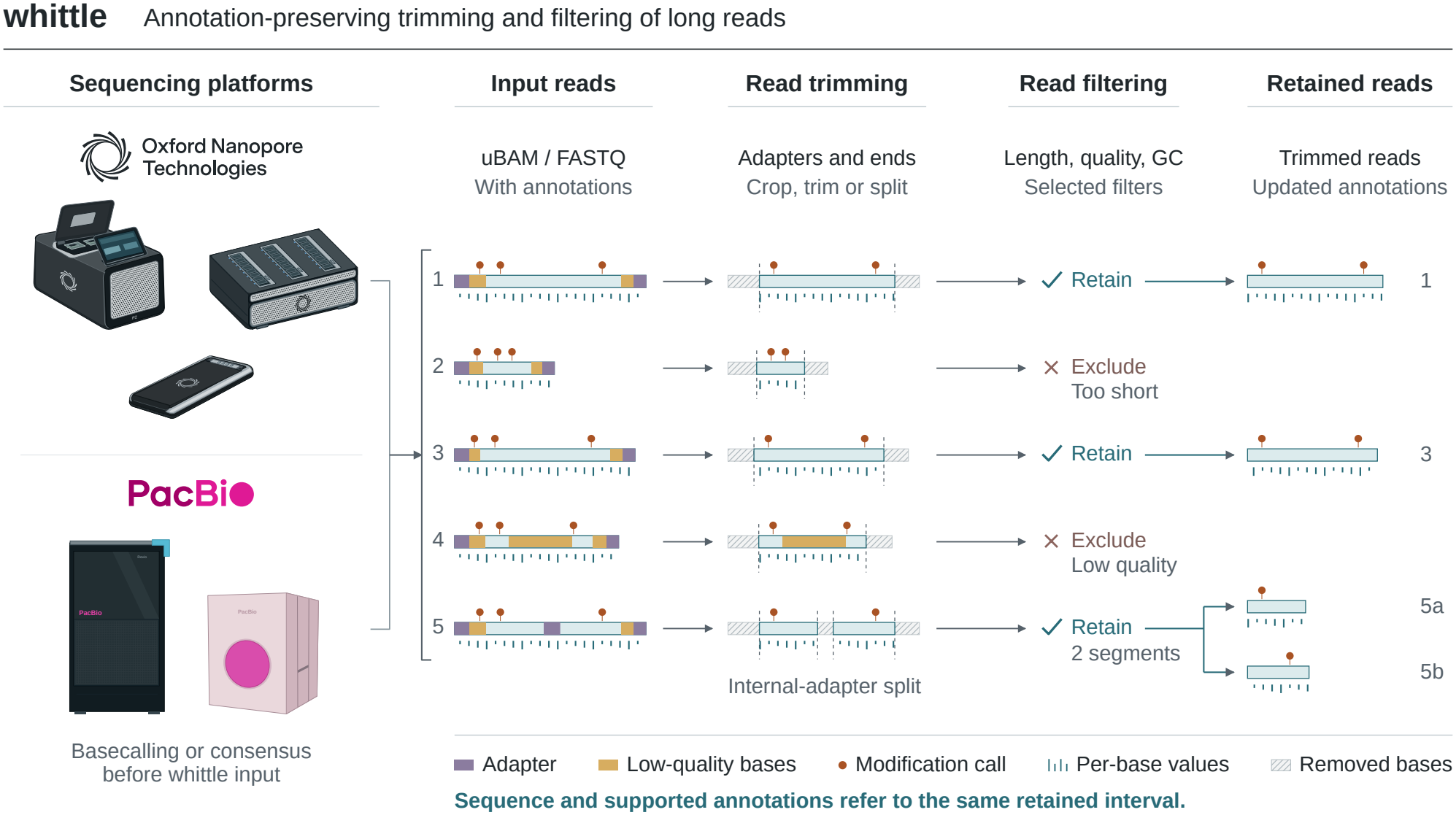

## 1 Introduction

Long reads from Oxford Nanopore Technologies (ONT) and PacBio instruments often retain adapters, barcodes and primers from library preparation. Chimeric reads can contain an adapter between two sample sequences (inserts). Preprocessing removes these technical sequences from read ends and splits chimeric reads to recover the inserts. Among existing tools, chopper filters and quality-trims reads (De Coster and Rademakers 2023, De Coster 2026). Beyond filtering, fastplong trims adapters and splits chimeras (Chen 2023, OpenGene 2026). Porechop_ABI discovers unknown adapters (Bonenfant et al. 2023), and cutadapt trims supplied sequences (Martin 2011).

ONT and PacBio basecalling software writes unaligned BAM (uBAM) records that can carry base-modification calls in `MM` (modification types and positions), `ML` (their probabilities) and `MN` (the sequence length they refer to; SAM/BAM Format Specification Working Group 2024). ONT records can also carry move tables linking bases to raw signal (Oxford Nanopore Technologies 2026a), and PacBio records can carry per-base polymerase kinetics (Pacific Biosciences 2023). `samtools fastq -T` copies selected tags into FASTQ headers, producing tagged FASTQ (Danecek et al. 2021). Trimming and splitting must remove the annotations of discarded bases and shift the rest to each output segment. Keeping the original tags leaves per-base arrays longer than the sequence and modification calls on removed bases. Dropping them discards information needed for modification analysis and signal-aware variant calling (Yu et al. 2026).

chopper 0.14.1 rewrites modification tags in tagged FASTQ but leaves other position-dependent tags unchanged (De Coster 2026). ONT’s basecaller, dorado, splits chimeric reads by default, and dorado trim updates modification and move-table tags when removing ONT adapters and primers (Oxford Nanopore Technologies 2026a). modkit repair restores modification tags on trimmed reads from the untrimmed records, but not on split reads (Oxford Nanopore Technologies 2026b). lima, PacBio’s demultiplexing tool for high-fidelity (HiFi) reads, clips modification and kinetic tags when it removes barcodes (Pacific Biosciences 2026a). We developed whittle to combine adapter removal, chimera splitting and quality filtering for both platforms in uBAM and tagged FASTQ, updating the supported annotations of every output segment in one pass over basecalled reads.

## 2 Features and implementation

### 2.1 Input, output and processing steps

From files or standard input, whittle reads uBAM and plain, gzip or BGZF FASTQ, tagged or untagged. uBAM records can also be exported as tagged FASTQ, optionally gzip-compressed. Aligned BAM is refused because trimming would invalidate the alignment.

Optional processing steps execute in a fixed order (Figure 1A; Supplementary Figure S1). A tag filter first selects reads by SAM tag values. Adapter processing removes terminal technical sequence, including verified barcodes at positions recorded by dorado, and splits reads at chimera junctions. Fixed cropping then removes a specified number of bases from each end. Quality processing can trim low-quality ends or split at low-quality runs. It can also retain the highest-scoring segment or all maximal-scoring segments (Ruzzo and Tompa 1999) under modified Mott scoring, which rewards bases above a quality cutoff. Each resulting segment is filtered by length, mean quality, expected errors (the sum of error probabilities; Edgar and Flyvbjerg 2015) and GC content.

**Figure 1.**
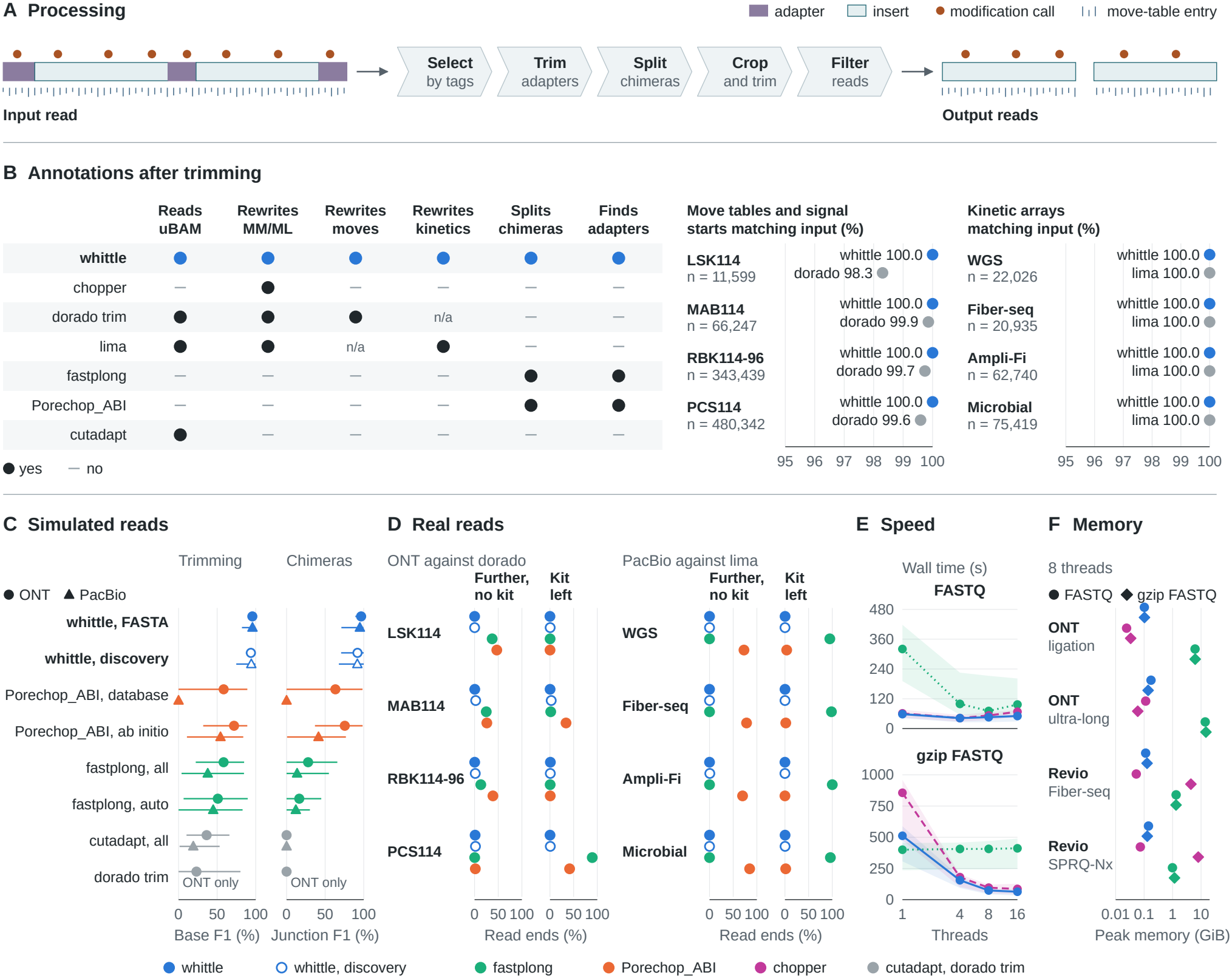
Processing and benchmarks (Supplementary Section S4). (A) Trimming and splitting an annotated read. (B) Tool capabilities (Supplementary Table S9) and annotation checks against values calculated from the untrimmed input, in reads that whittle and the reference tool (dorado for ONT, lima for PacBio) trimmed to the same bases. In the capability grid, n/a marks an annotation of the other platform. dorado’s records come from a separate basecalling run, and their match rates are lower bounds (Supplementary Section S4.4). Agreement within three bases, rather than exact agreement, is reported in Supplementary Table S4. (C) Means (markers) and ranges (lines) over eight simulated designs, including designs outside the scope of dorado trim and of Porechop_ABI’s adapter database (Supplementary Tables S2 and S3). dorado trim processes only ONT reads. The “whittle, FASTA”, “fastplong, all” and “cutadapt, all” runs received every design sequence, and “auto” marks automatic mode. (D) Read ends where a tool trimmed further than the reference without a kit match or past the last kit match (“Further, no kit”, possible loss of sample sequence) or kept kit sequence that the reference removed (“Kit left”, incomplete trimming). Lower values are better for both. Filled whittle markers show the kit preset (ONT) or kit FASTA (PacBio), and fastplong ran in automatic mode and Porechop_ABI in ab initio mode. (E) Wall time for cropping and filtering untagged FASTQ, as the median (lines) and range (bands) of four dataset values, where each value is the median of two runs. chopper’s lines are dashed and fastplong’s dotted. (F) Peak memory of the eight-thread runs, as two-run medians. For chopper’s gzip pipeline, memory is the larger peak of chopper and pigz.

### 2.2 Adapters, barcodes and primers

Adapters, barcodes and primers can be given as ONT and PacBio presets or FASTA files for other kits and custom libraries (Supplementary Section S3, Figure S3). Matching uses Sassy’s vectorised approximate search (Beeloo and Groot Koerkamp 2026), accepts IUPAC codes and handles sequences truncated at read ends. Stricter rules for internal matches limit chance splits and distinguish primer sites from chimera junctions (Supplementary Section S3.1).

When sequences are unknown, discovery assembles frequent 16-mers from the ends of up to 40,000 sampled reads and searches inward from each identified sequence for further flanks, barcodes and primers. In amplicon libraries, it uses variation between templates to separate primers from the conserved sample sequence that follows them. Discovered sequences can be reviewed before trimming (Supplementary Section S3.2).

### 2.3 Annotations

whittle updates each output segment’s annotations from its retained input bases (Figure 1B; Supplementary Figure S2, Table S1). It removes modification calls outside that segment, preserves remaining call probabilities (`ML`) and recalculates call positions (`MM` skip counts) and sequence length (`MN`). Invalid `MM`/`ML`/`MN` blocks are removed and counted.

Per-base arrays are sliced in their stored order. Reverse-strand PacBio kinetics are stored last base first, so whittle counts the retained interval from the other end of these arrays (Supplementary Section S2). PacBio query coordinates and read names are updated for the retained segment. fiber-tools adds nucleosome and accessibility tags to Fiber-seq reads, in which m6A marks accessible DNA (Stergachis et al. 2020, Jha et al. 2024). whittle removes these tags from trimmed or split records, because fibertools can recompute them. Most other tags are copied unchanged.

With `--update-moves` and uBAM output, whittle slices ONT move tables and adjusts the signal-start offset. Following dorado’s convention, split segments receive the parent read identifier and, with move updating, their offset in the parent signal. Without move updating, move-table tags are removed from trimmed records.

### 2.4 Implementation

whittle is written in Rust, using noodles for BAM, seq_io for FASTQ and libdeflate for compression. Reads are processed in parallel batches, optionally preserving input order. The x86-64 release binaries require AVX2.

## 3 Results

We compared whittle 1.0.0 with fastplong 0.7.1, Porechop_ABI 0.5.1, cutadapt 5.2, chopper 0.14.1 and dorado trim 2.1.2. dorado 2.1.0 basecalled, trimmed and demul-tiplexed ONT reads, and lima 26.2.1 trimmed and demulti-plexed PacBio reads.

### 3.1 Simulated reads

We simulated ONT and PacBio reads with Badread 0.4.2 (Wick 2019) at its default 95% mean identity, which is below PacBio HiFi accuracy. We recorded which bases came from adapters, barcodes and primers. Seven designs tested these sequences alone, in pairs or together. An eighth used sequences absent from whittle’s presets. Primer designs used whole 16S rRNA amplicons, and a control carried only Badread’s default terminal adapters. Each design had two replicates of 17,479 to 19,303 reads with a 5% chimera probability. Base-level scoring counted removed technical bases as true positives, retained technical bases as false negatives and removed insert bases as false positives.

With all sequences of each design supplied as FASTA, whittle reached mean base-level F1 scores of 95.6% for ONT and 95.9% for PacBio (Figure 1C; Supplementary Table S2). Given the same sequences, fastplong scored 58.4% and 37.8%, and cutadapt 36.4% and 19.0%. Both identified most technical bases but also removed insert sequence at matches inside reads (Supplementary Section S4.2). Without supplied sequences, whittle discovery reached 93.7% and 94.2%, Porechop_ABI ab initio mode 71.9% and 54.4%, and fastplong automatic mode 50.9% and 44.9%.

We counted a junction carrying technical sequence as recovered if both inserts were kept in separate output segments, each at least 90% complete. Splits elsewhere counted as false positives. With supplied sequences, whittle’s mean junction F1 was 96.6% for ONT and 95.0% for PacBio (Figure 1C). cutadapt, which does not split reads, recovered no junctions (Supplementary Table S3). In the ONT control, whittle discovery removed 0.10 insert bases per end, Porechop_ABI ab initio mode 4.79 and fastplong automatic mode 7.19. whittle discovery split a mean of one read per replicate and fastp-long automatic mode 2,145.

### 3.2 Real reads compared with dorado and lima

Four public ONT libraries covered human genomic DNA (SQK-LSK114), 16S mock communities (SQK-MAB114-24), rapid barcoding (SQK-RBK114-96) and cDNA (SQK-PCS114). We basecalled each library twice from the same signal, with and without dorado trimming (and demulti-plexing for barcoded libraries). All tools processed the untrimmed reads, including some already split by dorado (Supplementary Section S4.3). Trim positions within three bases of dorado’s counted as agreement. For larger differences, we searched the bases between the two trim positions for kit sequences (for ONT, those of whittle’s presets). We counted trimming beyond dorado as removing kit sequence only when kit matches reached the tool’s trim position. With insert boundaries unknown, we treated a kit match as evidence of technical sequence.

On the human library, whittle’s preset agreed with dorado at 91.95% of read ends and trimmed further with a kit match at another 7.71% (Supplementary Table S4). It trimmed further without a kit match at 0.21% (Figure 1D), mostly removing adapters with many sequencing errors. All 54 gaps between its split segments contained a kit match. fastplong automatic mode trimmed further without a kit match at 37.17% of ends, partly because it trims 10 bases past each adapter by default.

In the 16S library, whittle’s preset removed 16S primers left by dorado and trimmed further without a kit match at 0.41% of ends, against 24.71% for fastplong automatic mode and 25.86% for Porechop_ABI ab initio mode.

On the rapid-barcoding library, whittle’s preset agreed with dorado at 94.30% of read ends and trimmed further without a kit match at 0.32%. dorado assigned a barcode to 5% of these reads. On them, agreement fell to 52.42%, mostly because whittle trimmed further with a kit match (24.74% of ends), dorado trimmed further without a kit match (14.44%) or whittle kept kit sequence that dorado removed (8.02%; Supplementary Table S4). On the cDNA library, the preset agreed at 90.73% of ends and trimmed further without a kit match at 1.18%. Other tools agreed at 4.69% to 5.51% of cDNA ends.

lima’s trimming served as the reference for four PacBio Revio libraries with kinetics: human whole-genome (WGS), Fiber-seq, Ampli-Fi (amplified low-input) metagenome and microbial isolate. whittle received lima’s kit sequences or, for Ampli-Fi, the sequences most often clipped by lima. It agreed at 99.20% to 100.00% of read ends and never trimmed further without a kit match (Figure 1D). fastplong agreed at no more than 4.99%, and Porechop_ABI, which targets ONT reads, at no more than 29.99%. On four other PacBio datasets, discovery trimmed into conserved 16S sequence at 5.3% of amplicon ends and split 6.6% of cDNA reads at Alu repeats (Supplementary Section S4.3).

### 3.3 Annotations

We checked signal tags in 901,627 ONT records that whittle and dorado trimmed alike (to the same bases), including 712 split segments. In every record, whittle’s move tables and signal-start offsets matched values that a standalone script calculated from the untrimmed input (Figure 1B). dorado’s records, from a separate basecalling run, matched in 98.3% to 99.9% of records per library. We did not test move tables after quality processing, in reads with several kept segments or in RNA reads (Supplementary Section S4.4). In 181,120 records that whittle and lima trimmed alike in the four Revio libraries, all four kinetic arrays of both tools matched the input slices. whittle’s modification calls matched the input in all 1,403,991 records trimmed alike in the eight PacBio datasets (Supplementary Table S5).

In 20,000 reads from each speed-benchmark dataset (Section 3.4), we decoded modification calls with HTSlib (Bonfield et al. 2021). After adapter and quality processing, all 68,144,389 retained calls in 232,504 uBAM records and their tagged FASTQ counterparts matched the expected annotations (Supplementary Table S6).

### 3.4 Speed and memory

We benchmarked cropping and filtering on ONT ligation and ultra-long reads and on Revio Fiber-seq and SPRQ-Nx chemistry reads, 29.980 to 30.015 Gb each except Fiber-seq (16.941 Gb). We ran each configuration twice on a 24-core Intel Core Ultra 9 285K (Figure 1E). Tools cropped 10 bases per end and kept reads of at least 500 bases and mean quality Q10, with adapter search disabled.

Across the four datasets at eight threads, whittle processed untagged gzip FASTQ 5.1 to 5.8 times as fast as fastplong and 1.2 to 1.4 times as fast as chopper, using 54.4% to 61.3% of chopper’s CPU time (Supplementary Table S7). In these untagged gzip runs, whittle’s peak memory stayed at or below 142 MiB; fastplong’s reached 15,187 MiB on ultra-long reads (Figure 1F). chopper’s gzip pipeline used less memory than whittle on ONT reads but up to 8,151 MiB on PacBio reads.

whittle used 45.1% to 55.4% of chopper’s CPU time on tagged gzip FASTQ and processed the ligation uBAM directly in 102 seconds at eight threads. At one and four threads, partly cached input gave chopper the shorter median wall time on ligation plain FASTQ, but whittle used less CPU time (Supplementary Section S4.5).

## 4 Conclusion

whittle trims, splits and filters ONT and PacBio reads from uBAM or tagged FASTQ in one preprocessing step, updating the supported annotations of each output segment. Independent checks found no mismatches in the tested move tables, modification calls or PacBio kinetic arrays. At eight threads, whittle cropped and filtered gzip FASTQ in less wall and CPU time than chopper. Discovery can recover unknown technical sequences but can mistake conserved or repeated sample sequence for technical sequence in amplicon and cDNA libraries. We recommend kit presets or supplied FASTA sequences for these libraries.

## Data availability

The ONT libraries compared with dorado are on the ONT Open Data registry (Oxford Nanopore Technologies 2026c), and the PacBio libraries compared with lima come from the Human Pangenome Reference Consortium, PacBio and the European Nucleotide Archive (Human Pangenome Reference Consortium 2026, Pacific Biosciences 2026b, Swiss Institute of Bioinformatics 2026; Supplementary Section S4.3). The cropping and filtering datasets are ONT and Revio Fiber-seq uBAMs from the same consortium and PacBio’s Revio SPRQ-Nx Use 3 release (Human Pangenome Reference Consortium 2026, Pacific Biosciences 2026b; Supplementary Section S4.5). The simulated reads, the scripts that generated and scored them, and the commands of every benchmark are archived at https://doi.org/10.5281/zenodo.24051464 (Kiliç 2026).

## Author contributions

Erdi Kiliç (Conceptualization; Software; Validation; Formal analysis; Writing, original draft; Writing, review and editing). Burçak Otlu (Supervision; Writing, review and editing).

