## Supplementary Data for "whittle: annotation-preserving trimming and filtering of long reads"

### S1 Usage and processing order

The processing steps in whittle are optional and chosen via the command line, but they always execute in the order listed below. The step numbers match the row numbers of Figure S1A, where row 0 is the input read. Each step operates only on the sequence retained by the preceding steps (Figure S1).

1. The tag filter retains or discards entire reads according to their SAM tag values. It is applied prior to any trimming and before discovery selects its sample of reads.
2. Adapter processing removes adapters, barcodes and primers from the read ends and splits reads at chimera junctions (Section S3). Steps 3 to 6 treat each resulting piece, or a read without a junction, as a segment.
3. Barcode restriction removes each barcode that it confirms at a position recorded by dorado, together with the bases before a front barcode or after a rear barcode. This step is executed only if adapter processing is also selected.
4. Fixed cropping removes a specified number of bases from the front (`-H`, `--trim-front`) and tail (`-T`, `--trim-tail`) of each segment. Each segment is cropped only once, so the pieces produced by a subsequent quality split are not cropped again.
5. Quality processing trims or splits segments according to base quality.
6. The filters evaluate the length, GC content, mean quality and expected errors of each remaining segment.

The first example reads and writes uBAM, trims the LSK114 adapters, updates the move tables (Section S2) and drops output segments shorter than 500 bases (`-l 500`):

```
whittle -i reads.bam -o trimmed.bam --adapter-preset lsk114 --update-moves -l 500
```

For FASTQ output from uBAM input, whittle copies all SAM tags, or those specified with `--fastq-tags`, into the read headers and removes the move-table tags from trimmed records (Section S2). The second example writes gzip-compressed FASTQ with headers containing only the modification tags `MM`, `ML` and `MN`, updated for the retained bases. It trims the same adapters and keeps segments with a mean quality of at least Q10 (`-q 10`):

```
whittle -i reads.bam -o trimmed.fastq.gz --fastq-tags MM,ML,MN --adapter-preset lsk114 -q 10
```

The third example retains reads with a dorado quality score (`qs`) of at least 10 and a `dx` value of 0, which dorado assigns to simplex reads (single-strand reads) not used in a duplex read. Discovery draws its sample from these reads alone, and the adapters it finds are trimmed from them:

```
whittle -i reads.bam -o simplex.bam --tag-filter '[dx]==0 && [qs]>=10' --adapter-discover
```

Barcode restriction (Figure S1A, step 3) uses the front and rear barcode positions that dorado records in its `bi` tag when it demultiplexes reads (Oxford Nanopore Technologies 2026). whittle trims at a recorded position only if it detects a barcode within three bases of that position. By default, this search allows errors of up to 20% of the barcode length. The detected barcode may be either of two types: one of the barcodes that adapter processing searches for, or a built-in barcode with a number that corresponds to dorado's barcode call (`BC`). The run summary reports the number of reads in which whittle detects no barcode at a position recorded by dorado. This number includes reads that dorado has already trimmed.

`--quality-trim` selects one of four quality-trimming methods, and `--quality-cutoff` sets their Phred score threshold. This threshold has no default value and must be given with the method, because no single value suits all sequencing platforms and basecallers. Two of the methods, `best` and `segments`, rank sequence segments by a modified Mott score, which is the sum of the scores of their bases. The score of each base is the error probability at the cutoff minus the base's own error probability, with both values derived from Phred scores (Ewing and Green 1998). At a Q20 cutoff (error probability 0.01), a Q30 base raises the score by 0.009 and a Q10 base lowers it by 0.09.

- `ends` removes bases from each end of a segment until it reaches a base that meets or exceeds the quality cutoff. It therefore stops even at an isolated high-quality base within a low-quality tail.
- `best` retains the highest-scoring segment, which it identifies in a single pass. In contrast to `ends`, it trims through an isolated high-quality base within a low-quality tail, and the retained segment may still contain bases below the cutoff.
- `segments` keeps every maximal-scoring segment (Ruzzo and Tompa 1999) that scores at least as much as 50 error-free bases. No option changes this threshold. Two high-quality regions separated by a low-quality stretch remain distinct segments if the stretch lowers the score by more than the smaller of their two scores.
- `runs` splits a segment at every stretch of consecutive bases below the cutoff, provided that the stretch contains at least `--min-low-quality-run` bases (default 1). Shorter stretches remain in place, and below-cutoff bases are trimmed from both ends of each resulting piece.

Only `segments` and `runs` accept `--quality-end-cutoff`, which defaults to the value of `--quality-cutoff`. These two methods use it to trim the ends of each piece in the same way as `ends`. Raising this cutoff makes only the end trimming stricter; the split positions still follow the scoring rule of `segments` or the minimum run length of `runs`.

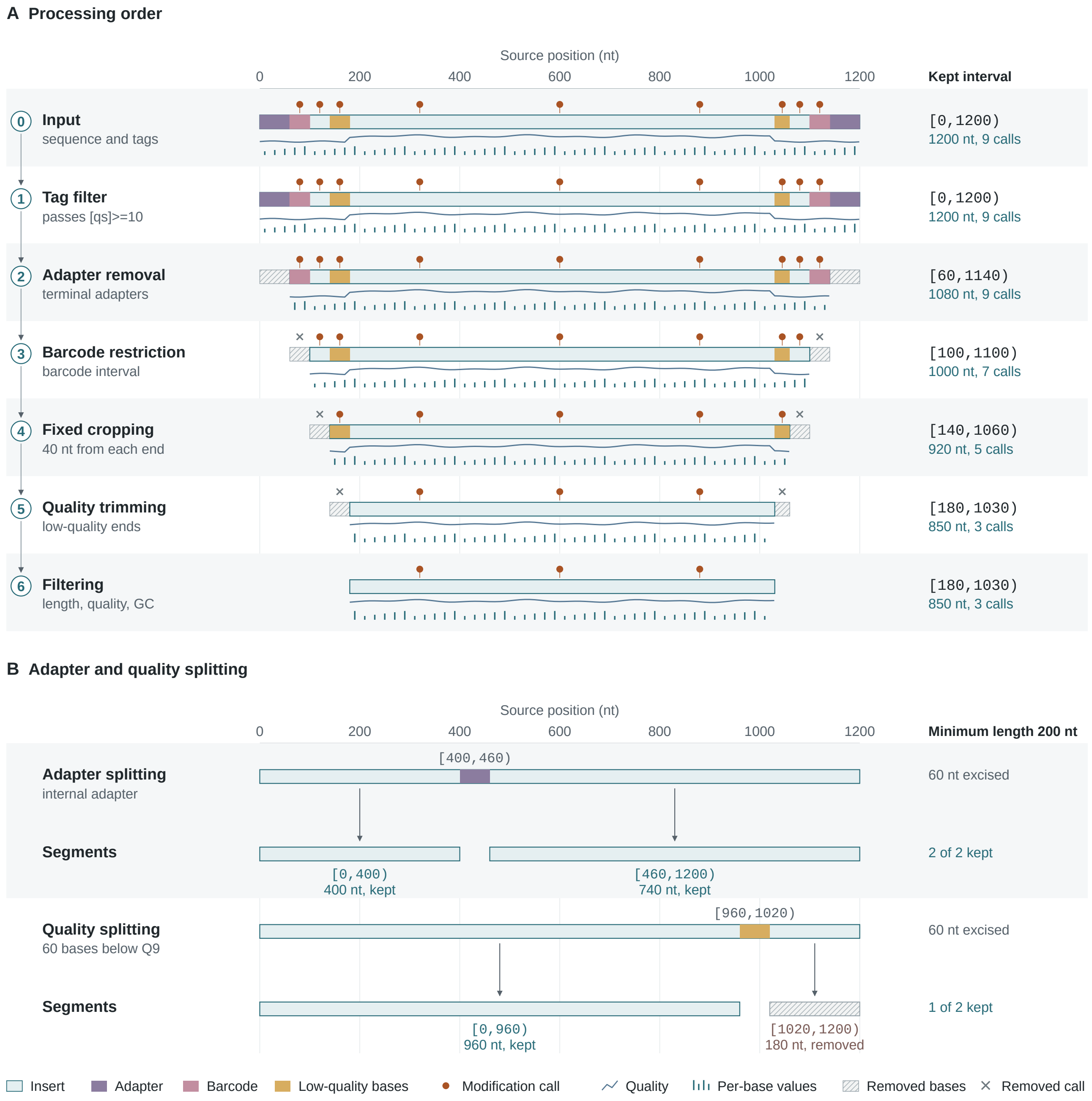


**Figure S1.** Processing order and read splitting. Positions and lengths are in nucleotides (nt), and intervals are half-open, including their start and excluding their end. (A) A read of 1,200 bases traced from the input (row 0) to filtering (row 6), with the kept interval, its length and the number of modification calls in it on the right. The read passes the tag filter `[qs]>=10`, and whittle confirms and removes the barcodes at the positions in dorado's `bi` tag (row 3). The kept interval narrows from [0,1200) to [180,1030), the calls outside it are dropped, and the read passes the final length, quality and GC filters. (B) Splitting at an internal adapter (upper rows) and at a run of 60 bases below Q9 (lower rows). The 180-base piece left by the quality split is shorter than the minimum output length of 200 bases and is discarded.

By default, the filters keep segments of at least 1 base (`-l`) with a mean quality of at least Q0 (`-q`). In the default quality mode, mean quality is the average per-base error probability of a segment, expressed as a Phred value. With `-q 10`, as in the second example, a segment passes if it has at most 0.1 expected errors per base, regardless of its length. `--quality-mode arithmetic` switches the filter to the arithmetic mean of the Phred scores, and `median` to their median. `--max-expected-errors` caps the total expected errors of a segment (Edgar and Flyvbjerg 2015), a limit suited to amplicons of similar length.

With `--rejected-output`, whittle writes rejected reads and segments to a separate file. This file has the format of the main output (uBAM or FASTQ) and is compressed according to its own file extension. Each rejected record carries a `wr` tag giving the reason for its rejection. `--summary-json` writes the run summary as a JSON report. The report gives the parameters of the run, the counts of reads, bases and rejected segments, and any annotation problems.

An ONT read that remains a single segment after the trimming steps retains its original name. If multiple segments are produced, each takes the suffix `_segment_N`, where N numbers the segments in read order, starting from 1. whittle assigns the numbers before the filters run, so a segment keeps its suffix even when it is the only one to pass. Suppose that read `r1` is split into three segments and the second is too short. In this case, `r1_segment_1` and `r1_segment_3` are written to the main output, and `r1_segment_2` is written to the rejected output with the `wr` tag `too_short`. If the input is uBAM or tagged FASTQ, the segments of a split PacBio read with a name that follows the PacBio conventions do not take the `_segment_N` suffix. Instead, the name of each segment ends in its own query coordinates, which give the start and end of the segment on the original read (Section S2). With untagged FASTQ input, every split read takes the `_segment_N` suffix.

With more than one thread, whittle writes each record as soon as the record is ready. If `--preserve-order` is given, whittle instead writes the records in their input order.

whittle stops with an error message that names the read when it encounters any of four kinds of input record:

1. An aligned record, since trimming would invalidate its alignment.
2. A uBAM record in which the base qualities do not cover every base.
3. An unaligned record with the reverse-complement flag, which indicates that its SEQ field holds the reverse complement of the read. whittle trims in read orientation, so it would cut the wrong ends of such a record and misplace its position-indexed tags.
4. A record with an `Mm` or `Ml` tag. These tags are the older names of `MM` and `ML` and were written by earlier basecallers. whittle rewrites only `MM` and `ML`, so the calls in `Mm` and `Ml` would refer to the wrong bases after trimming.

With `--update-moves` and uBAM output, the run also stops when a trimmed read carries a move table but its read group does not name a DNA or RNA basecalling model (Section S2). Other problems, such as an invalid modification block or a per-base array of the wrong length, are reported but do not stop the run.

### S2 Annotation handling

With positions counted from zero, a segment that keeps the bases s to e − 1 of an input read of L bases is written [s,e). The annotation value of each base in [s,e) moves from input position p to output position p − s, and the values of removed bases are discarded (Figure S2A and B). Adapter trimming, fixed cropping and quality processing all follow this rule. Tags without a row of their own in Table S1 are copied unchanged, with one exception: whittle adds the roles of the searched sequences (adapter, primer or barcode) to dorado's trim-mode field `tm`. A `tm` value that is not in dorado's format is left unchanged.

**Modification calls.** `MM` stores each call position as a skip count, the number of canonical bases (C, for example) passed over between one call and the next (SAM/BAM Format Specification Working Group 2024). whittle removes the calls that lie outside the segment and recalculates the skip counts of the remaining calls so that they count only the canonical bases within the retained sequence. It also sets `MN` to the length of the output read (Figure S2B). `ML` holds the probabilities of the remaining calls in their original order.

The `MM`, `ML` and `MN` tags of a record together form its modification block. whittle removes invalid blocks from the output and counts them. A block is invalid if `MM` cannot be parsed or if its skip counts place a call beyond the last base of the canonical type in the read. It is also invalid if `ML` has the wrong type or length, or if `MN` differs from the read length. A lone `ML` or `MN` tag in a record without a string-valued `MM` is copied unchanged.

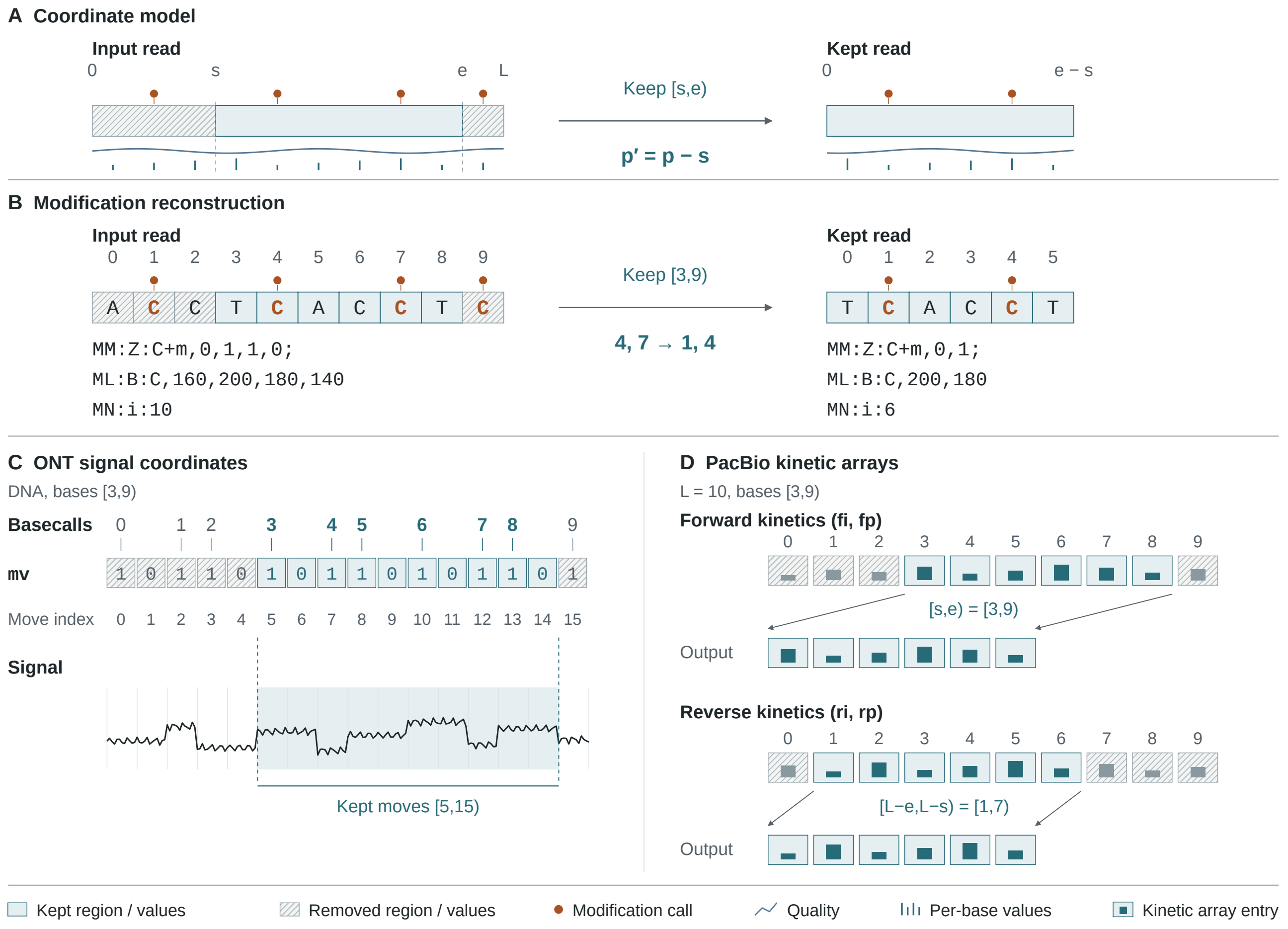


**Figure S2.** Annotation updates after a crop. Bases and array entries are numbered from zero, and L is the input read length, which is 10 bases in panels B to D. (A) Coordinate rule for a segment that keeps the bases [s,e): input position p becomes output position p′ = p − s, and e − s bases remain. (B) Modification tags of the read `ACCTCACCTC` cropped to [3,9), shown in SAM text format. The calls at positions 1 and 9 are dropped and those at 4 and 7 become calls at 1 and 4. `MM` changes from `C+m,0,1,1,0` to `C+m,0,1`, `ML` keeps only the probabilities 200 and 180, and `MN` falls from 10 to 6. (C) Move table (`mv`) of a DNA read, with one entry per block of signal samples. An entry of 1 marks a block in which a new base starts, and an entry of 0 marks one in which the current base continues. A base can span several blocks, so the 10 bases take 16 entries (0 to 15). The base numbers above the table mark the entry at which each base begins, and the move indices below it number the entries. Bases 3 to 8 occupy entries 5 to 14. The leading stride of `mv` is not drawn. (D) PacBio kinetic arrays, drawn in their stored order, which the output keeps. The forward arrays keep entries 3 to 8. The reverse arrays are stored last base first, with entry i holding the value of base L − 1 − i. They keep entries 1 to 6, the values of bases 8 down to 3. Kinetic values are illustrative.

A modification group is one entry of `MM`: a header such as `C+m`, followed by the skip counts of its calls. If all calls of a group fall outside the segment, the group remains in `MM` without skip counts. Such an empty group still carries information. In the default implicit mode of `MM`, bases of the group's canonical type that carry no call are interpreted as unmodified. An empty group therefore declares that every retained base of its canonical type is unmodified (every C of the output read, in the case of `C+m`). Removing the group would leave the state of these bases unknown.

**Per-base arrays.** Some BAM tags are arrays that hold one value for each base of the read. whittle recognises multiple such arrays by name, including the PacBio kinetics (Pacific Biosciences 2023). Among the kinetic arrays, `fi` and `fp` hold the inter-pulse durations and pulse widths of the forward strand, and `ri` and `rp` hold those of the reverse strand. whittle also treats any other BAM array as per-base if its length equals the read length. This applies to custom arrays as well, even when their values do not refer to single bases. Two kinds of array are exempt from this length test. Tags with a rule of their own, such as `ML` and `mv`, follow that rule instead. The PacBio arrays `sn`, `ac` and `bc` hold a fixed number of values regardless of the read length and are not sliced.

whittle slices each per-base array over [s,e). The reverse-strand kinetics `ri` and `rp` are the exception: they are stored last base first, so whittle slices them over [L−e,L−s) (Figure S2D). If the length of an array recognised by name differs from the read length, whittle retains the array unchanged and counts the read under `malformed_tag_reads` among the warnings of the run summary. This rule also applies to an empty array recognised by name, which therefore remains empty.

The run-length coverage tag `sa` describes PacBio subread coverage: each run of bases with equal coverage is written as its length followed by the coverage. When the run lengths add up to the read length, whittle crops `sa` to the kept bases and re-encodes it; otherwise it keeps the tag unchanged and counts the read in the same warning. For example, in a 10-base read, `4,5,3,7,3,5` gives coverage 5 to bases 0 to 3, coverage 7 to bases 4 to 6 and coverage 5 to bases 7 to 9. A crop to [2,7) keeps two bases at coverage 5 and three at coverage 7, so `sa` becomes `2,5,3,7`.

**PacBio query coordinates and names.** The tag `qs` has two meanings: an integer query coordinate in PacBio records and a float mean quality in ONT records. whittle distinguishes them by the type of the value. PacBio `qs` and `qe` give the start and end positions of the record on the original read (Pacific Biosciences 2023). whittle updates them only if a record carries both. It then sets output `qs` = input `qs` + s and output `qe` = input `qs` + e. With uBAM or tagged FASTQ input, whittle also updates read names that follow the PacBio conventions, such as `movie/1/ccs` and `movie/1/100_106`. If such a name ends in query coordinates, whittle rewrites it to show the kept interval. When a read with such a name is split, each segment is named after its own coordinates. The coordinates in the output name are counted from the input `qs` if the input record carries both `qs` and `qe`. Otherwise, they are counted from the start coordinate in the input name, or from 0 if that name has none. Split reads with other names take the `_segment_N` suffix (Section S1).

Cropping the first base from a six-base read named `movie/1/100_106` (`qs` 100, `qe` 106) gives `movie/1/101_106`, with `qs` 101 and `qe` 106. When a 10-base read named `m64011_190830_220126/123/ccs` (`qs` 100, `qe` 110) is split into [0,2) and [3,10), its segments are named `m64011_190830_220126/123/ccs/100_102` and `m64011_190830_220126/123/ccs/103_110`, each with `qs` and `qe` that agree with its name.

Whole-read PacBio tags such as `rq` (predicted read accuracy) and `np` (number of passes) are copied unchanged. `ds` and `ls` hold the data that skera and lima need to undo their processing (skera splits Kinnex arrays into segments). Both tags describe the untrimmed read, so whittle removes them (Table S1).

**ONT mean quality.** whittle recomputes the float `qs` of ONT records from all kept quality values, as the Phred value of their mean error probability. The result can differ from dorado's `qs` for the same bases, because in reads longer than 60 bases, dorado excludes the first 60 bases from its mean (Oxford Nanopore Technologies 2026).

**ONT signal tags.** The move table `mv` begins with the stride, which is the number of raw signal samples in each block. The stride is followed by one entry per block, in the 0/1 encoding drawn in Figure S2C (Oxford Nanopore Technologies 2026). The signal-start offset `ts` gives the sample at which the first block begins. In dorado records, the integer signal endpoint `ns`, the duration `du` and the start time `st` describe the signal of the whole read. The sample rate is `ns` / `du`. A read split from a parent read also carries `sp`, its offset in samples within the parent signal, and `pi`, the parent's identifier.

Move updating (`--update-moves`) rewrites the signal tags of trimmed records in uBAM output. It requires a read-group description that specifies a DNA or RNA basecalling model, because the model determines the signal direction (Section S1). In RNA reads, the signal runs in the opposite direction to the reported sequence, so whittle maps the kept interval to [L−e,L−s) before locating the move-table entries.

With move updating, the move table for each output record contains only the entries that correspond to its own bases. In the case of a record cropped from an unsplit read, `ts` advances by one stride for each removed entry that precedes the retained entries. The signal of a DNA read ends at its last base, and the signal of an RNA read ends at its first base. If a crop removes the base where the signal terminates, `ns` is set to the end of the last retained block, at output `ts` + stride × number of retained entries. Conversely, if the crop keeps this base, `ns` remains at its input value, since the move table can position the signal end only at a stride boundary.

For example, cropping the DNA read of Figure S2C to bases 3 to 8 keeps entries 5 to 14 and advances `ts` by five strides. Because the crop removes base 9, where the signal terminates, `ns` becomes the output `ts` plus ten strides. A crop leaves `sp` and `st` unchanged. However, whenever a crop changes `ns`, whittle rescales `du` by the same factor, so that `ns` / `du` still gives the sample rate.

With move updating, a split record receives a `ts` of 0 and an `ns` equal to the number of signal samples covered by its segment. Its `sp` is set to the offset of the segment within the parent signal. Following dorado's convention, whittle also recomputes `du` and `st` from the sample rate of the input read. The `du` of the record is its own `ns` / input sample rate, and its `st` is the input `st` advanced to the first signal sample of the segment. Both tags stay unchanged when the input has no positive `ns` or no positive float `du`. An `st` that cannot be read as a timestamp also stays unchanged.

whittle keeps the poly(A) estimates `pt` and `pa` only with move updating and only when the tail lies inside the kept signal. In a split record, it shifts `pa` into the record's own signal coordinates (Table S1).

Without move updating, as in any FASTQ output, whittle removes `mv`, `ts` and the integer `ns` from trimmed records. It also removes them if the move table is not a 0/1 table with exactly one 1 per base. In a split record, `sp`, `st` and a float `du` are removed as well, unless its signal tags are rewritten from a move table.

As dorado does when it splits a read (Oxford Nanopore Technologies 2026), whittle gives every ONT split record the parent identifier `pi` and a read number `rn`, with or without move updating. It also sets the MinKNOW event count `me` and end reason `er` in the same way, under the conditions in Table S1. whittle does not modify the raw signal files.

**Fiber-seq footprints.** fibertools stores the start and length of each nucleosome and accessible patch of a Fiber-seq read in four integer arrays: `ns`, `nl`, `as` and `al` (Jha et al. 2024; Fiber-seq 2026). Their read coordinates do not fit a trimmed or split read, and since fibertools can rebuild the arrays from the kept modification calls, whittle removes them from such records. whittle distinguishes the two `ns` tags by type: an array-valued `ns` is the fibertools tag, and an integer `ns` is dorado's signal endpoint.

**Table S1.** Annotation updates for a segment that keeps the bases [s,e) of an input read of L bases (Figure S2A). Updates assume valid input annotations, and `zmw/qs_qe` stands for any read name that ends in its query interval.

| Annotation | Update | Exceptions |
|---|---|---|
| `MM`, `ML`, `MN` | calls outside [s,e) removed, skip counts and length recomputed, probabilities of kept calls retained | invalid blocks are removed and counted. Standalone `ML` and `MN` are unchanged |
| Forward per-base arrays | sliced over [s,e) | a known array of the wrong length is kept, with a warning |
| PacBio reverse kinetics (`ri`, `rp`) | sliced over [L−e,L−s) | a known array of the wrong length is kept, with a warning |
| Run-length coverage (`sa`) | cropped and re-encoded | kept unchanged when the runs do not cover the read |
| PacBio query coordinates and names (`qs`, `qe`, `zmw/qs_qe`) | moved to the new interval | unpaired query-coordinate tags are unchanged. Names without an interval are unchanged for unsplit reads |
| ONT mean quality (`qs`, a float in dorado records) | recomputed from the kept qualities | none |
| ONT signal (`mv`, `ts`, `ns`, `sp`, `pi`, `du`, `st`) | sliced with `--update-moves`, otherwise `mv`, `ts` and the integer `ns` removed, and `sp`, `st` and float-valued `du` also removed from split records. `pi` is set on every split record | `mv`, `ts` and the integer `ns` removed for unsupported move encodings. Unknown model direction is an error |
| ONT poly(A) estimates (`pt`, `pa`) | kept and shifted with `--update-moves` when the tail is kept, otherwise removed | none |
| ONT read number, event count and end reason (`rn`, `me`, `er`) | set on split records as dorado sets them when splitting reads | `me` and `er` only when present, and `er` only on split records that do not keep the end of the parent signal |
| Barcode positions (`bi`), PacBio skera and lima tags (`ds`, `ls`) | removed, as they describe the untrimmed read | none |
| fibertools footprints (`ns`, `nl`, `as`, `al` arrays) | removed from trimmed or split records | none |
| Other tags | copied unchanged | dorado's trim mode `tm` (a read-group field in uBAM, a per-read tag in tagged FASTQ) gains the roles of the searched sequences (adapter, primer, barcode) |

### S3 Adapter matching and discovery

whittle finds technical sequences at read ends and inside reads by approximate matching. It obtains these sequences from kit presets or a FASTA file, or infers them from the reads by discovery (Section S3.2).

#### S3.1 Sequence matching

Each searched sequence has exactly one role: adapter, barcode or primer. The role determines whether a match inside a read can split the read and, if so, under which conditions. Every preset entry has a predefined role. The ONT presets cover kits for ligation, rapid and ultra-long sequencing; for native, rapid and rapid PCR barcoding; for cDNA-PCR and direct RNA sequencing; and for 16S and ITS amplicons. For PacBio, a single preset covers SMRTbell libraries. A FASTA file supplies the sequences of other kits or of a custom library. The role of a FASTA entry is set by its header description, the text after the entry's name. If the description contains the whole word `primer` or `barcode`, whichever of the two words occurs first sets the role. An entry with neither word is an adapter.

whittle narrows a kit preset to the sequences present in a sample of the reads, so that sequences absent from the library neither slow the search nor cause spurious trims. If a kit preset is the only source of sequences, whittle searches for every preset sequence in up to 2,000 reads from the start of the input. This number of reads is the default sample size of `--adapter-sample-reads` without discovery. A preset sequence is retained if it trims or splits at least three of these reads and at least 0.2% of them. If no sequence reaches both counts, whittle keeps the whole preset. Narrowing applies only to presets. whittle searches for every sequence of a supplied FASTA file, because narrowing on a sample could remove a rare custom adapter. Adding a FASTA file to a preset turns narrowing off. When discovery runs with a preset, whittle narrows only the preset's barcode entries, using the discovery sample (Section S3.2).

**Matches at read ends.** whittle searches the w bases at each end of every read (the search range, 150 bases by default) for sequences of all three roles, on both strands. The approximate search runs in Sassy (Beeloo and Groot Koerkamp 2026), which uses SIMD (single instruction, multiple data) operations. A match may contain errors up to the error allowance of its sequence. This allowance is sequence length × error rate (20% by default), rounded down. If the read end cuts a sequence short, so that only part of it is present in the read, whittle still removes it through a partial match to that part (Figure S3A). A partial match must cover at least 10 bases. Its error allowance equals that of a sequence nine bases shorter than the part it covers. At the default error rate of 20%, a partial match of 10 to 13 bases must therefore be exact. An IUPAC code in a searched sequence matches any of the bases it stands for (Figure S3B).

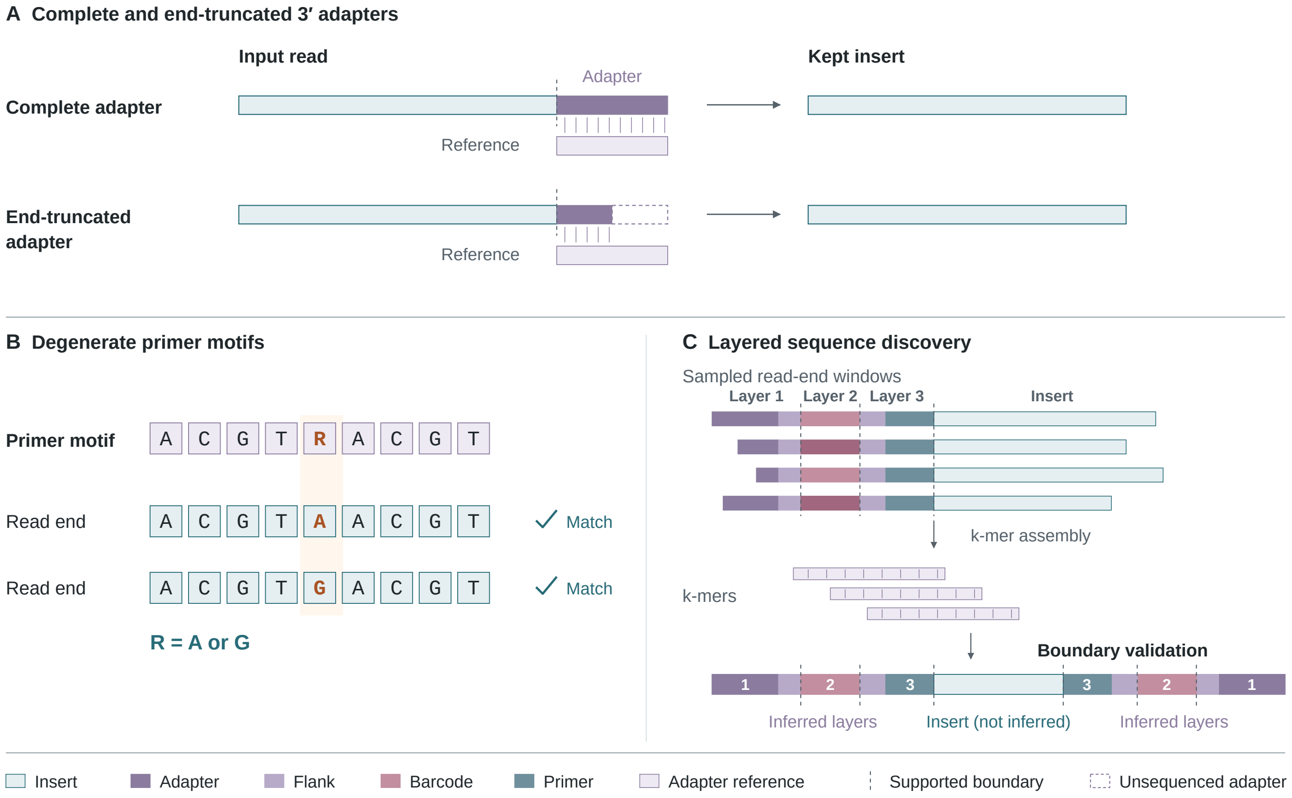


**Figure S3.** Adapter matching and discovery. (A) A complete 3′ adapter and one truncated at the read end, both matched against the adapter reference and removed, leaving the insert. Ticks mark matched bases, and a dashed box marks the adapter bases that were not sequenced. (B) A degenerate primer, in which the IUPAC code R accepts A or G. (C) Layer-by-layer discovery from the read ends inwards. Read-end k-mers (16-mers in whittle) are assembled into candidate sequences, their boundaries are validated, and the inferred adapter, flank, barcode and primer layers are separated from sample sequence. The read-end windows, one per sampled read end, lack varying numbers of the adapter's outermost bases and carry one of two barcodes, drawn in two shades. The validated read is shown with both ends. Its layers are numbered 1 to 3 from each end towards the insert: the adapter and its flank (1), the barcode (2) and the flank and primer next to the insert (3). Supported boundaries are drawn as dashed vertical lines, separating the insert from the adapter in A and adjacent layers in C. An inferred primer can keep sample bases when there is too little variation near it (Section S3.2). Sequences are illustrative.

**Matches inside reads.** whittle evaluates a match inside a read more strictly than a match at a read end, so that fewer reads are split at chance matches or at primer sites in sample sequence. Inside a read, only a match to the whole sequence counts, and the role of the sequence determines when such a match splits the read:

- An adapter splits a read of any length.
- A single primer or barcode splits a read only if its expected number of exact chance matches in a read of that length is at most $10^{-4}$ (see Chance matches, below).
- A 16S or ITS primer also matches inside genomic reads that span an rRNA operon. Such a primer therefore splits a read only if it lies next to a junction partner. This partner is either another technical sequence removed at the same junction or a second such primer in the orientation expected at a junction. In an amplicon library, however, the primer splits a read on its own (see Amplicon libraries, below).
- Where the constant flanks on either side of a barcode can identify the junction, the flanks, not the barcode, split the read.
- The UMI (unique molecular identifier) pattern in PCS114 does not split reads, and the primer beside it marks the junction instead.
- With MAB114 as the sole preset, whittle assumes an amplicon library. The 16S and ITS primers of the preset then take the adapter role, and each internal match to them marks a chimera junction, regardless of read length.

**Amplicon libraries.** Unless MAB114 is the sole preset, whittle treats a library as an amplicon library only if the sampled reads meet two conditions:

- At least 60% of the sampled reads carry a primer at one or both ends, immediately behind any other technical sequence at that end.
- For each sampled read, whittle calculates the share of sampled reads with lengths within 5% of the length of this read. The median of these shares is at least 15%.

whittle measures both shares on at most 4,000 reads spread evenly over the sample. A 16S or ITS primer counts towards the first share without further conditions. Any other primer counts only if the inserts behind it start alike. whittle evaluates this condition on the read ends at which the primer borders the insert. At least 3% of these read ends, and at least 10 of them, must share one 8-base word starting at any of the first five positions of the insert. Words in which a single base occupies three quarters or more of the positions, as in a poly(A) tail, do not count. As in discovery (Section S3.2), these thresholds are fixed.

**Chance matches.** A longer read offers more positions at which a match can occur by chance, so whittle allows fewer errors in the internal matches of longer reads. To estimate how often chance matches occur, whittle models read bases as independent and uniformly distributed. For a sequence, q(k) is the probability of a chance match with at most k errors (substitutions, insertions or deletions) at a given start position. If a base of the sequence is an IUPAC code representing d bases (d = 1 for A, C, G and T), it matches a random read base with probability d/4.

whittle obtains q(k) by summing the probabilities of alignment paths. If multiple alignments can produce the same match, the sum counts that match more than once. The sum therefore overstates the true probability, and the allowances derived from it are conservative. A read of L bases offers each sequence N = 2L start positions on its two strands, so the expected number of chance matches of the sequence in the read is N × q(k).

whittle assigns each read to a length class. The upper limits of the classes are 4,096, 8,192, 16,384 bases and so on, each limit double the one before, and every read of a class is shorter than its limit. Internal allowances are computed per class, with L at the class limit. Within a class, a sequence that can split reads receives the highest k at which N × q(k) is at most $10^{-4}$, but never more than its error allowance. If even the exact chance matches of a sequence exceed this bound in a class, an adapter splits the reads of that class at exact matches only (k = 0). In that case, a primer or barcode does not split them.

For each sequence, the expected number of inexact chance matches (those with at least one error) is N × (q(k) − q(0)). The sum of these numbers over all sequences that can split reads must also be at most $10^{-4}$ per read. While the sum exceeds this bound, whittle lowers the allowance of the sequence with the largest N × q(k), and of every sequence tied with it, by one error and then recalculates the sum. The sum counts a sequence and its reverse complement as one term and each barcode of a panel of alternative barcodes as a separate term. If two 16S or ITS primers meet at a junction, whittle matches them with a pair allowance of their own. whittle sets this allowance in the same way, so that the expected number of chance pairs is at most $10^{-4}$ per read. A chance pair consists of two chance matches of such primers, the second starting no more than 11 bases before or after the end of the first. No sequence's allowance ever increases in a longer class.

For example, the probability that a 16-base barcode without IUPAC codes matches exactly by chance at a given start position is q(0) = $4^{-16}$. For reads of 65,536 to 131,071 bases, L is set to the class limit of 131,072 bases. The expected number of exact chance matches, N × q(0), is then 6.1 × $10^{-5}$, which is within the bound. In the next longer class, N doubles and this number rises to 1.2 × $10^{-4}$, which exceeds the bound. The barcode therefore cannot split a read of 131,072 bases or more.

**Allowances at read ends.** The same model caps error allowances at read ends. At each read end, a match can start at any of w + 1 positions, 0 to w bases from the end. Each sequence therefore has N = 4(w + 1) start positions over both ends and both strands. Two bounds apply. First, a per-sequence cap allows each sequence no more than 0.1 expected chance matches per read over these positions. Second, the expected number of inexact chance matches, N × (q(k) − q(0)), summed over all sequences, must be at most 0.1 per read. whittle applies this second bound twice, each time lowering the allowances of the largest contributors one error at a time, as for internal matches. Each sequence receives two allowances in this way. The anchored allowance covers matches that start within 11 bases of the read end or of a match already accepted at that end, and it is computed over 4 × 12 start positions per sequence. The range-wide allowance covers matches anywhere in the search range and is computed over all 4(w + 1) positions. It is never larger than the anchored allowance. A match with more errors than its range-wide allowance is trimmed only once it is anchored.

At the default error rate of 20% and w = 150, the per-sequence cap lowers the allowance at read ends by one error for sequences of 11, 12 and 15 bases without IUPAC codes. Without the cap, their allowances of two, two and three errors would give expected chance matches of 0.55, 0.16 and 0.15 per read. Sequences of 13 and 14 bases keep an allowance of two errors, because their additional bases lower their expected chance matches to 0.048 and 0.014.

#### S3.2 Sequence discovery

Discovery infers the adapters, barcodes and primers of a library from sequence that recurs at the ends of many reads, without a kit catalogue. It proceeds from each read end inwards, one layer of technical sequence at a time (Figure S3C). The outermost layer, such as an adapter at the physical end, is found before any flank, barcode or primer behind it.

The discovery sample is taken from the start of the input and holds 40,000 reads by default (`--adapter-sample-reads`). Sampling stops earlier if the sample reaches 256 MiB of record data or 67.1 million bases. Only two of the settings in this section can be changed with command-line options: the sample size and the error rate, which determines the error allowances (Section S3.1). All other thresholds, limits and window sizes in this section are fixed in the code and were not tuned on the benchmark datasets of Section S4.

At each read end, discovery keeps a search boundary from which it looks for the next layer. The boundary initially lies at the physical read end. If a kit preset or FASTA file is also supplied, the boundary instead starts behind the matches of the supplied sequences. It moves inwards past every layer found in the read.

**Step 1: count.** whittle examines a window of 100 bases at each read end, extending inwards from the search boundary. It then counts how many of these windows contain each 16-mer.

**Step 2: assemble.** whittle links the 500 most frequent 16-mers into candidate sequences through overlaps of 15 bases. The counts along a candidate show where a layer ends: they stay steady over sequence shared by the reads and drop at the start of sample sequence or of a barcode panel. They rise where a barcode, present in only some of the reads, meets a flank that all of them carry. For example, if every read has the same adapter followed by one of 12 equally common barcodes, the counts drop to about one twelfth where the barcodes start. A candidate ends where its counts leave one stable level for another, subject to four rules:

- (a) The two levels differ at least fourfold. A degenerate primer base with two equally common alternatives only halves the counts of the 16-mers that span it, a change too small to meet this rule.
- (b) Before the shift, the counts remain at one level over at least eight consecutive 16-mers. A gradual climb with no stable level, as when reads lack varying numbers of an adapter's outermost bases, does not end a candidate.
- (c) Four 16-mers between the levels, such as those spanning a barcode and its flank, may have intermediate counts.
- (d) The second level persists across 24 consecutive 16-mers. If fewer remain before the candidate's inner end, it must persist across all of them, and at least four must remain. A single sequencing error lowers the counts of at most 16 consecutive 16-mers, while a barcode alters the counts over a longer run.

**Step 3: test.** whittle accepts a candidate only if it is frequent near the search boundary and rare in read interiors. To measure its frequency near the boundary, whittle searches for the candidate, within the error allowance, in at most 4,000 windows that extend up to 200 bases inwards from the boundary. The candidate must occur in at least 1% of these windows and in at least 20 of them. This absolute minimum limits sparse discoveries in small samples.

The candidate must also be anchored at the boundary, with at least 60% of its matches starting within 50 bases of the boundary. The margin allows for remnants of the previous layer, which may be partly eroded or vary in length. A majority suffices because layers further out may have gone undetected in some reads, and in these reads the candidate starts further from the boundary.

whittle evaluates rarity only in reads with at least 400 bases between their two search boundaries. The interior windows for this test are 100 bases long and lie 100 to 200 bases inwards from each boundary. In these windows, the candidate must occur at less than one quarter of its rate at the read ends.

**Step 4: move inwards.** In each read, if a match to an accepted candidate begins no more than 50 bases from the search boundary, whittle advances that boundary to the inner end of the match.

**Forms at the two ends.** Because whittle searches every sequence on both strands (Section S3.1), one discovered sequence can serve both read ends: it occurs unchanged at one end and as its reverse complement at the other. In two cases, discovery also stores the form at the other end as a sequence of its own.

- Unequal lengths: as with the two strands of a Y-shaped adapter, the forms at the two ends differ in length and end at different bases towards the insert. Storing both forms allows an internal copy of either form to match without the bases it lacks counting as errors.
- Same orientation: a sequence such as a hairpin adapter can appear in the same orientation at both ends. In this case, trimming with the sequence accepted at one end leaves part of the form at the other end in place. Discovery keeps that form as a layer of the other end and continues behind it.

**Degenerate layers.** A UMI or similar degenerate sequence yields no frequent 16-mers, so whittle finds it from the base composition, position by position, behind each accepted layer and towards the read centre. It skips barcode layers, because the sequence expected behind a barcode is a flank shared by the reads rather than a degenerate sequence.

At each position, whittle compares the frequency of each base with p, the share of that base in the outermost 300 bases of both read ends, over the sampled reads. A base predominates at a position if its frequency reaches (1 + p)/2, the midpoint between p and 1. A base is depleted at a position if its frequency is p/4 or lower. A fixed technical base, read at the platform's per-base accuracy, is expected to reach the midpoint. The lower threshold targets tags designed so that their variable positions leave out a base. At a position in a genomic insert, each base is expected to occur at about its share, which lies between the two thresholds whatever the composition. A degenerate layer requires at least 16 consecutive positions, each with one predominant base or at least one depleted base.

Discovery also tests the variable positions, those at which no base predominates, for randomness. Across the read ends, no combination of the bases at these positions may occur 20 or more times. The number of distinct combinations must also be at least half the number of read ends. Layers formed by a barcode panel, by conserved sample sequence or by the primers of the two strands show only a few combinations and fail.

A degenerate layer is written with IUPAC codes, like the degenerate positions of a primer. For instance, a random tag with A, C or G (code V) at its variable positions and fixed T spacers between them is written as VVVVTTVVVVTTVVVVTTVVVVTTT. The layer takes the primer role, so the primer rules of Section S3.1 decide whether it splits reads.

**Barcodes between flanks.** Discovery recovers barcodes that sit between two constant flanks by clustering the stretches that the flanks enclose. It applies this clustering to a layer only if two conditions hold. First, the median start of the most common candidate of the layer lies at least 11 bases inside the boundary. Second, this candidate is at least four times as common as any candidate with a median start fewer than 11 bases inside the boundary. Discovery then assumes that the stretch between the boundary and this candidate holds one barcode per read.

Each window holding this candidate contributes the stretch from the boundary to the match, unless the stretch is shorter than 11 bases, the minimum length of a searched sequence. Clustering proceeds one cluster at a time. Each cluster starts from a seed, the most common stretch that is still unassigned, and gathers every unassigned stretch that contains this seed within the error allowance. If a cluster has at least 20 members and these make up 0.1% or more of the windows, the consensus of its members becomes a barcode. No layer yields more than 128 barcodes.

**Roles and names.** Without preset or FASTA sequences, a candidate of layer 1 becomes an adapter when its median distance from the physical read end is no more than 20 bases. However, if such a candidate matches a 16S or ITS primer, it does not take the adapter role, and only the primer rules of Section S3.1 then determine whether it splits reads. Every other discovered sequence, deeper layers included, is a barcode if it forms part of a barcode panel and a primer otherwise.

whittle names the discovered sequences `inferred_1`, `inferred_2` and so on. Each is shown beside the built-in or supplied sequence that matches it best, if their identity is at least 85%. A lower threshold would let a discovered sequence of 16 to 32 bases match an unrelated entry by chance, because it is compared with the whole catalogue in both orientations. With `--adapter-report`, whittle prints the inferred sequences and trims nothing.

**Sample sequence inside a candidate.** A candidate may extend from an adapter or primer into conserved sample sequence, such as a gene start shared by many reads. Discovery separates the two by comparing their depths (distances from the physical read end) at the two read ends. When library preparation gives both ends of a molecule the same layers, technical sequence reappears as its reverse complement at the opposite read end, at about the same depth as at its own end. Conserved sample sequence can also occur at the opposite read end, but it need not follow the same layers there, so its depth can differ.

Depths are medians over the reads, measured from the physical read end to the sample-facing edge of each match. D denotes the depth of the candidate at its own read end. whittle divides the candidate into overlapping 16-base pieces that start four bases apart and scans these pieces outwards, beginning at the sample-facing end of the candidate. If the sample-facing edge of a piece lies x bases further out than that of the candidate, the depth of the piece at its own end is $D - x$.

For each piece, whittle searches for its reverse complement in the 300 bases nearest the opposite read end. The search uses an error allowance computed from half the error rate, so that a match does not extend past the bases present at the read end. The reverse complement counts as found if it occurs in at least 20 reads and in at least one eighth as many reads as the candidate. The one-eighth threshold is low because a sequence from one end of the molecule, such as a conserved gene start, appears at the opposite read end only in reads of the other strand. The stricter search also misses some of these copies. $D'$ denotes the depth of a found reverse complement at the opposite read end.

Each piece has one of three outcomes. If $D'$ and $D - x$ differ by more than 35 bases, the piece counts as sample sequence. whittle then trims the candidate to end at the piece's outer edge, $D - x - 16$ bases from the read end, and continues the scan. If they differ by 35 bases or fewer, the piece marks technical sequence and the scan stops. A piece leaves the candidate unchanged if its reverse complement is not found, and the scan then continues.

Suppose that the reads begin with a 30-base adapter, a 20-base primer and 40 bases of conserved sample sequence. The opposite end of these reads is truncated: it finishes with the reverse complement of the conserved stretch, followed by the reverse complement of only 7 primer bases. For the innermost piece ($x = 0$), $D = 90$ and $D' = 47$, a gap of 43 bases, so the piece is cut away as sample sequence. Each piece further out shows the same gap and is cut away in turn, up to the piece that reaches 8 bases into the primer. At the opposite end, the reverse complement of this piece lacks one of its 16 bases. It is still found, because at half the default error rate, the error allowance of a 16-base piece is one error. The cuts remove all of the conserved sequence, leaving the adapter and the 12 primer bases next to it as the inferred sequence.

**Primer boundaries in amplicon libraries.** Assembly can join a primer and the conserved gene start that follows it into a single candidate. Reads of the other strand that reach the far end of the molecule also cover this primer. In these reads, the reverse complements of the primer and the gene start recur together at the opposite read end, so the depth check cannot separate them. Discovery relies instead on variation between templates to identify the template start, where the primer ends and the sequence copied from the template begins. It places this start at the first variable position near the primer where the reads carrying the two most common bases at that position also differ from each other deeper in the insert.

The method rests on two assumptions. First, the primer of a read, including any degenerate bases, carries no information about the template from which the read was copied, in contrast to the sequence that follows the primer. Second, every variant of a conserved start belongs to a group of related templates, and these groups diverge deeper in the insert.

A position marks the template start only if it meets three conditions:

1. Two read groups: among the reads aligned to the candidate, those with the most common base at the position form one group and those with the second most common base form the other. The smaller group must hold at least 20 windows and at least 5% of the aligned windows.
2. Different templates: deeper in the insert, each group carries 12-mers of its own. whittle takes the 12-mers from the 100 bases that begin 16 bases beyond the candidate's inner end. A 12-mer counts as a group's own if 20 or more windows of the two groups contain it and 90% or more of these windows belong to that group. In each group, at least a quarter of the windows must contain such a 12-mer. Variation caused by recurrent sequencing errors or degenerate primer bases is expected in the reads of every template and therefore fails this condition.

3. Similar surroundings: past the position, the two groups mostly carry the same bases. Among the next 16 positions, whittle considers those read by at least 10 windows in each group. At least two thirds of these positions must show the same most common base in both groups. A sample variant within a conserved region should pass. Alternative primers, for instance the primers of the two strands, and panel barcodes differ at many nearby positions and should fail.

In the read windows aligned to the candidate, the evaluation begins 16 bases before the outer end of the match. It then advances inwards one position at a time and ends at the first base past the inner end of the match. The earliest position to meet all three conditions marks the template start. Discovery trims the candidate to end just before it, or discards the candidate if fewer than 11 bases remain. It tests the two ends of the gene on separate sets of reads.

An inferred primer can still carry conserved sample bases that lie before the first informative variant. We recommend kit presets or FASTA sequences for libraries made from a single template or with little variation near the primer.

### S4 Benchmarks

The benchmark archive (Kılıç 2026) holds the simulated reads, their truth tables and the scripts that generated them in `data/simulated/`. The scripts that download and prepare the real datasets of Section S4.3 are in `data/real/`. Each benchmark has its own folder, which holds its commands, its scoring and validation scripts and its raw results. Section S4.2 draws on `benchmark/adapter-accuracy/`, Sections S4.3 and S4.4 on `benchmark/dorado-reference/` and `benchmark/lima-reference/`, and Sections S4.4 and S4.5 on `benchmark/real-data/`.

#### S4.1 Software and hardware

All benchmarks used the same binary of whittle 1.0.0 (archived at https://doi.org/10.5281/zenodo.24051287), compiled in release mode with Rust 1.98.1 and the flag `-C target-cpu=x86-64-v3`. We recorded its checksum before any benchmark ran.

- Compared tools: fastplong 0.7.1, Porechop_ABI 0.5.1 (installed from Bioconda), cutadapt 5.2, chopper 0.14.1, and dorado trim 2.1.2 on the simulated reads.
- Reference tools: dorado 2.1.0, which basecalled, trimmed and demultiplexed the real ONT reads, and lima 26.2.1, which trimmed and demultiplexed the PacBio reads.
- Cropping and filtering benchmark: Sequali 1.0.2 (Vorderman 2025), seqkit 2.14.0 (Shen et al. 2024), pigz 2.8 and GNU time 1.10.
- Read simulation: Badread 0.4.2.
- Other software: pysam 0.24.1 and samtools 1.24.

The main text cites the compared and reference tools, Badread and samtools.

dorado basecalled the real ONT reads on an NVIDIA GeForce RTX 4070 SUPER GPU. The comparisons on simulated reads, and those with dorado and lima on real reads, ran on an 8-core AMD Ryzen 7 5800H system with 30 GiB of memory.

For the cropping and filtering benchmark, we used a machine with 125 GiB of memory and a 24-core Intel Core Ultra 9 285K (8 performance and 16 efficiency cores). The machine ran Ubuntu 24.04.5 LTS with Linux kernel 7.0.0 and used the powersave CPU frequency scaling governor. The benchmark files were read from and written to an ext4 volume on NVMe storage.

Apart from its kit-specific settings and sequences, each tool configuration kept the same options on every dataset. In every benchmark, a thread count is the number of threads requested from a tool. We set no CPU affinity, so on the 285K any requested thread could run on a performance or an efficiency core.

#### S4.2 Simulated reads

To score trimming base by base and chimera splitting junction by junction, we simulated ONT and PacBio reads in a control and in eight designs, which differ in the technical sequences their reads carry. A wrapper that we wrote around Badread 0.4.2 placed the adapters, barcodes and primers of each design in the reads. The wrapper also recorded the origin of every output base in the truth tables of `data/simulated/`. If no design is given, the wrapper operates in Badread's standard fragment mode and produces the same reads as unmodified Badread run with the same seed.

ONT reads were simulated with Badread's `nanopore2023` error and quality models, and PacBio reads with its `pacbio2021` models. On both platforms, glitches, junk reads and random reads were disabled, and read identities were drawn from Badread's default distribution (`--identity 95,99,2.5`), with a mean of 95% and a maximum of 99%. At these identities, the PacBio reads were less accurate than HiFi reads.

**Designs.** Seven designs covered every combination of adapters (A), barcodes (B) and primers (P): each alone, the pairs AB, AP and BP, and all three with flanks (ABP). The ONT designs used Badread's default ligation adapter (SQK-LSK109), ONT's 24 native barcodes and the 16S primers 27F and 1492R. The PacBio designs used the SMRTbell adapter, 24 random barcodes of 16 bases and the same two primers. The eighth design, N, used sequences found in none of whittle's presets: random adapters, a random flank and a panel of 12 random barcodes. In the control, the only technical sequences on either platform were Badread's default ligation adapters at the read ends.

In the control and in designs A, B, AB and N, each insert was cut from the genome of *Escherichia coli* K-12 MG1655 (GCF_000005845.2; University of Wisconsin 2013). Fragment lengths followed a gamma distribution (`--length 6000,5000`) with a mean of 6,000 bases and a standard deviation of 5,000 bases. In the four primer designs (P, AP, BP and ABP), each insert was a whole 16S rRNA amplicon between the 27F and 1492R primer sites, taken from SILVA 138.2 (Quast et al. 2013).

The amplicons represent 1,000 bacterial genera and have a median length of 1,448 bases. In the designs without primers, a template was a genome fragment. In the primer designs, a template was the whole amplicon with its two primers. In BP and ABP, the template also carried the barcodes of the design and, in ABP, its flanks. Each read held one template or, if it was chimeric, two or more templates. Each template was read in either orientation with equal probability.

In every design and in the control, technical sequences were present at 95% of 5′ ends and 70% of 3′ ends. On average, these sequences retained 85% and 70% of their bases, respectively (`--start_adapter 95,85`, `--end_adapter 70,70`). Truncation treated the technical sequences at a read end as a single block and shortened this block from the outside inwards. A barcode behind a 5′ adapter, for example, lost bases only after the whole adapter was gone.

The chimera probability was 5%, and chimeric reads joined complete templates. As in Badread, a junction could hold two ligated sequences, each with a probability of 25%: one on the 3′ end of the preceding template and one on the 5′ end of the next. In designs A, B, AB and N, about 56% of junctions held neither ligated sequence and carried no technical sequence. In the control, no junction carried technical sequence. In the primer designs, the 25% probability applied to adapters alone: every junction held the primers of both templates, always with the barcodes and flanks of the design beside them. Every design and the control had two replicates, simulated with seeds 11 and 23. With a target of 20,000 reads per replicate, we set Badread's quantity (the total number of bases to simulate) to 20,000 × mean insert length. The replicates held 17,479 to 19,303 reads.

**Scoring.** Base-level scoring compared each tool's output with the recorded origin of every base. True positives (TP) are removed technical bases, false negatives (FN) are kept technical bases and false positives (FP) are removed insert bases; kept insert bases are not counted. The base-level F1 is 2TP / (2TP + FP + FN). Table S2 gives the mean of the two separately scored replicates for each design, and its Mean F1 column gives the mean over the eight designs, with equal weight for each design. Figure 1C and the main text report this mean. The tables omit precision and recall, which the archived score files give for every run.

We scored a junction only if it carried technical sequence. A scored junction is recovered when its two flanking inserts end up in different output segments, each keeping at least 90% of its bases within one segment. Recovered junctions count as true positives, the remaining scored junctions as false negatives and excess segments (defined below) as false positives.

A scored junction is resolved when no output segment spans it. Removing a whole insert beside a junction resolves the junction without recovering it. Each read is allowed one output segment plus one for every resolved junction, and any further segment is excess. If K output segments are produced from a read with R resolved junctions, the read therefore holds max(0, K − 1 − R) excess segments. Splitting an insert, or splitting at a junction without technical sequence, adds one to K and nothing to R. For example, if a tool resolves a read's only junction and also splits one of its inserts at a chance match, three segments result (K = 3, R = 1). One of these segments is excess. Junction F1 applies the base-level F1 formula to these counts, while the base-level scores measure where a trim or split falls.

Control reads have no junction that can be scored. For these reads, Table S2 gives the mean number of insert bases removed per insert end bordering an adapter or a junction, and Table S3 gives the number of reads split.

**Tool configurations.** We ran each tool in one or more configurations, listed below under their row names in Tables S2 and S3.

- whittle: "FASTA" (every design sequence in a FASTA file, `--adapter-fasta`), "preset" (a kit preset alone, `--adapter-preset`), "discovery" (discovery alone, `--adapter-discover`) and "preset and discovery" (both together).
- Porechop_ABI: "database" (its built-in database) and "ab initio" (ab initio mode, `-abi`).
- fastplong: "all" (the FASTA file given to whittle, `-a`), "outermost" (the outermost sequence at each read end, given with `-s` at the start and `-e` at the end) and "auto" (automatic mode).
- cutadapt: "all" (every design sequence, with `-g` for 5′ sequences and `-a` for 3′ sequences, and up to one removal round per simulated layer, `-n`) and "outermost" (the outermost sequence at each read end, likewise with `-g` and `-a`, and the default single removal round).
- dorado trim: the kit that matches the design (`-k`), on ONT reads only.

In its preset configurations, whittle used `pacbio` on PacBio reads. On ONT reads, it used `lsk114` in A, `nbd114` in B and AB, `mab114` in the primer designs and `all` (the union of every ONT preset and the PacBio preset) in N and the control.

No FASTA header labelled a sequence as a primer or a barcode, so whittle had the same information as fastplong and cutadapt. whittle treated every supplied sequence as an adapter, and a match to an adapter inside a read can split the read (Section S3.1).

In B and BP, where a barcode panel forms the outermost layer, the outermost configurations received the first barcode of that panel. fastplong's outermost configuration could not run on design P, because its `-s` option rejects the IUPAC codes of the outermost primer.

We requested four threads from every tool. whittle, fastplong and cutadapt had a minimum output length of 100 bases, and dorado trim had none. Porechop_ABI ran with its default setting, which discards split pieces shorter than 1,000 bases. cutadapt used an error rate of 0.2. Every fastplong run had quality filtering disabled (`-Q`) and used its default 10-base trimming extension beyond each adapter.

Sequences given to cutadapt with `-g` or `-a`, and to fastplong in a FASTA file, can match anywhere in a read. fastplong split a read at such an internal match, and cutadapt removed everything from the match to the read end. When supplied with all design sequences, the two tools removed most technical bases but also much insert sequence at these internal matches. Over the eight designs, both tools had a mean base recall of 83% to 89% and a mean precision of 12% to 50% on both platforms.

On PacBio reads, fastplong ran on seven designs in both its "all" and "outermost" configurations, and the outermost sequences gave the higher F1 in six. The gain was marginal in B and largest in AB (91.0% against 16.5%; Table S2).

**Time and memory.** GNU time recorded the wall time and peak resident memory of every tool invocation (Table S2). We ran the invocations one at a time on the AMD system (Section S4.1), with plain FASTQ as input and output. Timings include writing the output but exclude the scoring. They also exclude a separate `--adapter-report` run of whittle, which only printed the inferred sequences (Section S3.2). The configurations differ in the work they do: discovery first infers the sequences from sampled reads.

**Table S2.** Trimming accuracy, time and memory on simulated reads. F1 (%) is the base-level trimming F1, Wall (s) the wall time and RAM (MiB) the peak resident memory. Mean F1 (%) is the mean F1 over the eight designs, as reported in the main text and Figure 1C. If a configuration has no F1 value in one of the designs, its mean is n/a. Every value is the mean of two replicates run with four requested threads. Designs carry adapters (A), barcodes (B), primers (P), pairs of these (AB, AP and BP), all three with flanks (ABP) or sequences found in no preset (N). The control reads carry only Badread's default ligation adapters. For these reads, the Control column gives the average number of insert bases removed by a tool per insert end that borders an adapter or a junction. Rows give the tool and its configuration (Section S4.2). Porechop_ABI's built-in database ("database") holds only ONT adapters and barcodes, and dorado trim leaves barcodes and 16S primers in the reads (dorado removes barcodes during demultiplexing). In designs outside their scope, these two configurations score 0.0. n/a marks a configuration that could not run or that needed design sequences the control lacks. For each platform, bold marks the highest mean F1 and, in each design, the highest F1 and the lowest wall time, memory and control value, together with any value tied with them at the displayed precision.

*ONT*

| Tool | Mean F1 (%) | A | | | B | | | P | | | AB | | | AP | | | BP | | | ABP | | | N | | | Control |
|---|---|---|---|---|---|---|---|---|---|---|---|---|---|---|---|---|---|---|---|---|---|---|---|---|---|---|
| | | F1 (%) | Wall (s) | RAM (MiB) | F1 (%) | Wall (s) | RAM (MiB) | F1 (%) | Wall (s) | RAM (MiB) | F1 (%) | Wall (s) | RAM (MiB) | F1 (%) | Wall (s) | RAM (MiB) | F1 (%) | Wall (s) | RAM (MiB) | F1 (%) | Wall (s) | RAM (MiB) | F1 (%) | Wall (s) | RAM (MiB) | |
| whittle, FASTA | 95.6 | 94.2 | **0.21** | **31.3** | 90.1 | 0.53 | **32.7** | **90.9** | **0.08** | **12.7** | **97.8** | 0.54 | **34.7** | **98.5** | **0.10** | **12.8** | **98.5** | 0.38 | **14.1** | 98.7 | 0.45 | **14.3** | 96.1 | **0.31** | **34.4** | n/a |
| whittle, preset | 82.8 | **95.3** | 0.31 | 54.1 | 89.1 | **0.26** | 58.0 | 87.9 | 0.11 | 19.8 | 97.3 | **0.39** | 55.9 | 96.9 | 0.19 | 19.4 | 95.2 | **0.09** | 19.8 | 98.4 | **0.16** | 20.8 | 2.4 | 0.49 | 56.4 | 0.91 |
| whittle, discovery | 93.7 | 94.7 | 1.35 | 224.2 | 91.4 | 2.34 | 215.1 | 89.7 | 4.65 | 130.6 | 97.7 | 3.64 | 224.0 | 96.3 | 2.09 | 128.8 | 87.2 | 6.85 | 133.5 | 96.1 | 7.14 | 130.7 | **96.6** | 3.56 | 223.9 | **0.10** |
| whittle, preset and discovery | **95.8** | 95.2 | 1.65 | 210.4 | **91.7** | 3.98 | 217.0 | 90.5 | 2.50 | 88.0 | 97.3 | 2.26 | 212.3 | 98.2 | 2.71 | 87.1 | 97.9 | 2.50 | 89.9 | **99.2** | 2.59 | 88.1 | 96.2 | 6.60 | 225.7 | 1.06 |
| Porechop_ABI, database | 58.4 | 83.5 | 126.20 | 772.2 | 83.3 | 541.29 | 772.3 | 0.0 | 53.77 | 512.4 | 89.2 | 902.30 | 772.4 | 72.8 | 62.81 | 511.9 | 65.0 | 166.26 | 511.6 | 73.6 | 326.64 | 509.5 | 0.0 | 90.02 | 772.0 | 4.77 |
| Porechop_ABI, ab initio | 71.9 | 83.3 | 279.86 | 778.6 | 83.2 | 646.73 | 778.6 | 32.0 | 179.21 | 518.5 | 89.1 | 1,058.57 | 778.7 | 61.8 | 299.03 | 517.9 | 59.4 | 316.51 | 517.9 | 84.1 | 497.78 | 518.8 | 82.5 | 548.51 | 778.1 | 4.79 |
| fastplong, all | 58.4 | 70.1 | 5.04 | 587.4 | 54.5 | 55.96 | 542.1 | 22.3 | 3.20 | 146.2 | 74.1 | 56.76 | 548.6 | 68.9 | 3.84 | 145.7 | 67.1 | 43.17 | 148.0 | 85.0 | 49.58 | 141.9 | 24.9 | 33.04 | 429.7 | n/a |
| fastplong, outermost | n/a | 74.3 | 5.77 | 409.5 | 6.9 | 5.73 | 398.5 | n/a | n/a | n/a | 65.8 | 2.31 | 401.6 | 69.7 | 0.95 | 100.2 | 4.7 | 1.52 | 107.9 | 30.7 | 1.35 | 103.7 | 69.5 | 5.81 | 401.9 | n/a |
| fastplong, auto | 50.9 | 71.5 | 5.97 | 587.7 | 6.5 | 6.62 | 558.7 | 15.6 | 1.19 | 139.7 | 64.1 | 2.48 | 577.9 | 58.1 | 0.85 | 140.2 | 20.6 | 0.91 | 142.5 | 89.6 | 0.98 | 149.1 | 81.1 | 6.45 | 580.3 | 7.19 |
| cutadapt, all | 36.4 | 30.2 | 6.01 | 38.8 | 25.7 | 125.57 | 41.1 | 26.3 | 1.25 | 35.5 | 36.6 | 258.35 | 40.7 | 48.3 | 3.96 | 33.8 | 47.6 | 81.79 | 37.5 | 65.9 | 198.68 | 37.0 | 10.3 | 53.35 | 35.6 | n/a |
| cutadapt, outermost | 22.4 | 33.7 | 5.42 | 39.1 | 7.9 | 4.35 | 39.4 | 20.9 | 1.03 | 37.2 | 22.1 | 2.96 | 39.4 | 40.3 | 0.88 | 36.0 | 5.8 | 0.93 | 37.3 | 21.3 | 1.27 | 36.8 | 26.9 | 5.73 | 39.4 | n/a |
| dorado trim | 23.2 | 80.2 | 2.56 | 554.0 | 0.0 | 2.23 | 617.1 | 0.0 | 1.36 | 515.7 | 42.8 | 0.98 | 610.0 | 45.5 | 0.69 | 541.5 | 0.0 | 0.77 | 532.4 | 17.2 | 0.82 | 537.8 | 0.0 | 1.92 | 621.0 | 0.28 |

*PacBio*

| Tool | Mean F1 (%) | A | | | B | | | P | | | AB | | | AP | | | BP | | | ABP | | | N | | | Control |
|---|---|---|---|---|---|---|---|---|---|---|---|---|---|---|---|---|---|---|---|---|---|---|---|---|---|---|
| | | F1 (%) | Wall (s) | RAM (MiB) | F1 (%) | Wall (s) | RAM (MiB) | F1 (%) | Wall (s) | RAM (MiB) | F1 (%) | Wall (s) | RAM (MiB) | F1 (%) | Wall (s) | RAM (MiB) | F1 (%) | Wall (s) | RAM (MiB) | F1 (%) | Wall (s) | RAM (MiB) | F1 (%) | Wall (s) | RAM (MiB) | |
| whittle, FASTA | **95.9** | 98.7 | 0.26 | **30.7** | **82.2** | 0.44 | **33.7** | **94.1** | 0.08 | **12.8** | **98.3** | 0.66 | **33.9** | **99.6** | 0.14 | **12.8** | 97.6 | 0.40 | **13.9** | 99.5 | 0.46 | **13.8** | 96.8 | 0.32 | **28.5** | n/a |
| whittle, preset | 36.7 | 98.7 | **0.24** | 53.0 | 0.5 | **0.22** | 54.1 | 0.0 | **0.07** | 19.2 | 79.1 | **0.26** | 54.4 | 73.2 | **0.13** | 19.2 | 0.0 | **0.08** | 19.1 | 42.0 | **0.11** | 20.3 | 0.2 | **0.23** | 53.7 | **0.00** |
| whittle, discovery | 94.2 | **98.9** | 2.94 | 226.2 | 74.8 | 1.29 | 215.7 | 91.5 | 3.12 | 125.7 | 98.2 | 7.31 | 225.2 | 96.4 | 5.33 | 121.2 | **97.8** | 3.31 | 119.3 | **99.7** | 11.73 | 116.0 | 96.5 | 3.19 | 225.0 | 0.11 |
| whittle, preset and discovery | 94.5 | 98.7 | 1.15 | 200.3 | 74.8 | 1.64 | 213.3 | 91.5 | 3.49 | 122.7 | 97.7 | 2.93 | 230.4 | 99.5 | 3.35 | 113.2 | 97.1 | 3.72 | 112.4 | 99.5 | 3.66 | 116.5 | **97.1** | 3.61 | 220.2 | 0.11 |
| Porechop_ABI, database | 0.0 | 0.0 | 90.63 | 772.4 | 0.0 | 81.46 | 772.6 | 0.0 | 53.24 | 512.4 | 0.0 | 70.62 | 772.1 | 0.0 | 54.48 | 511.4 | 0.0 | 56.66 | 512.4 | 0.0 | 53.24 | 509.7 | 0.0 | 74.01 | 772.4 | 5.44 |
| Porechop_ABI, ab initio | 54.4 | 78.7 | 554.07 | 778.7 | 10.9 | 358.19 | 778.7 | 31.8 | 248.88 | 518.7 | 64.8 | 468.28 | 778.3 | 48.5 | 260.78 | 517.4 | 39.6 | 268.80 | 519.0 | 76.8 | 236.12 | 515.7 | 83.9 | 509.56 | 778.6 | 5.45 |
| fastplong, all | 37.8 | 77.6 | 3.68 | 506.8 | 3.9 | 90.14 | 278.6 | 33.8 | 3.15 | 152.4 | 16.5 | 101.48 | 401.0 | 84.8 | 3.88 | 146.3 | 17.5 | 40.34 | 133.0 | 42.4 | 43.50 | 120.9 | 26.2 | 33.61 | 432.5 | n/a |
| fastplong, outermost | n/a | 85.7 | 5.16 | 383.4 | 3.9 | 4.72 | 356.5 | n/a | n/a | n/a | 91.0 | 4.50 | 412.4 | 85.1 | 0.94 | 102.9 | 6.7 | 1.00 | 110.5 | 50.8 | 0.80 | 101.2 | 70.4 | 5.87 | 383.3 | n/a |
| fastplong, auto | 44.9 | 68.8 | 5.64 | 521.0 | 0.5 | 5.09 | 404.4 | 27.2 | 0.88 | 146.5 | 63.7 | 4.84 | 518.5 | 83.0 | 0.96 | 147.5 | 37.0 | 1.12 | 146.3 | 0.0 | 0.84 | 132.0 | 79.4 | 6.03 | 580.0 | 7.26 |
| cutadapt, all | 19.0 | 25.5 | 7.80 | 36.7 | 1.3 | 89.05 | 34.9 | 27.0 | 1.34 | 38.2 | 5.0 | 190.39 | 36.7 | 53.2 | 5.52 | 35.4 | 8.2 | 46.64 | 31.4 | 22.6 | 102.69 | 31.9 | 9.5 | 62.06 | 38.0 | n/a |
| cutadapt, outermost | 21.0 | 20.7 | 7.72 | 37.2 | 0.6 | 4.10 | 35.3 | 21.2 | 0.82 | 35.7 | 20.6 | 7.30 | 38.4 | 42.5 | 1.20 | 34.7 | 4.7 | 0.72 | 34.1 | 31.4 | 1.23 | 36.7 | 26.2 | 5.67 | 37.9 | n/a |

**Table S3.** Chimera splitting on simulated reads. F1 (%) is the junction F1, and Mean F1 (%) is its mean over the eight designs, as reported in the main text and Figure 1C. The False splits column gives the number of excess output segments (Section S4.2). Both kept (%) is the percentage of scored junctions (those with technical sequence) that were recovered, with each of the two inserts keeping at least 90% of its bases in its own output segment. Every value is the mean of two replicates run with four requested threads. The Control column gives the mean number of reads split in the control design. Each replicate of the control holds approximately 19,100 reads and no junction with technical sequence. Design labels, row labels and n/a are as in Table S2, which also gives the wall time and peak memory of these runs. For each platform, bold marks the highest mean F1 and, in each design, the highest F1 and Both kept values, the lowest control value and any value tied with them at the displayed precision.

*ONT*

| Tool | Mean F1 (%) | A | | | B | | | P | | | AB | | | AP | | | BP | | | ABP | | | N | | | Control |
|---|---|---|---|---|---|---|---|---|---|---|---|---|---|---|---|---|---|---|---|---|---|---|---|---|---|---|
| | | F1 (%) | Both kept (%) | False splits | F1 (%) | Both kept (%) | False splits | F1 (%) | Both kept (%) | False splits | F1 (%) | Both kept (%) | False splits | F1 (%) | Both kept (%) | False splits | F1 (%) | Both kept (%) | False splits | F1 (%) | Both kept (%) | False splits | F1 (%) | Both kept (%) | False splits | |
| whittle, FASTA | **96.6** | 95.4 | 91.2 | 0 | 89.2 | 80.5 | 0 | **96.7** | **93.6** | 0 | **97.9** | **96.0** | 1 | **98.2** | **96.6** | 0 | **99.7** | **99.3** | 0 | 99.8 | **99.7** | 0 | **96.2** | 93.2 | 3 | n/a |
| whittle, preset | 84.0 | **95.5** | **91.3** | 0 | 89.2 | 80.5 | 0 | 96.6 | 93.5 | 0 | 96.0 | 92.6 | 1 | **98.2** | 96.4 | 0 | 97.6 | 95.3 | 0 | 99.3 | 98.6 | 0 | 0.0 | 0.0 | 1 | 1 |
| whittle, discovery | 91.9 | 93.4 | 87.7 | 0 | 84.5 | 73.2 | 1 | 96.5 | 93.2 | 0 | 97.2 | 94.5 | 0 | 70.7 | 55.6 | 0 | 99.3 | 98.7 | 0 | **99.9** | **99.7** | 0 | 93.7 | 91.3 | 15 | 1 |
| whittle, preset and discovery | 95.6 | 94.3 | 89.2 | 0 | **89.6** | **81.4** | 1 | **96.7** | **93.6** | 0 | 96.0 | 92.6 | 1 | **98.2** | **96.6** | 0 | 97.6 | 95.2 | 0 | 99.3 | 98.6 | 0 | 93.3 | 90.7 | 16 | 1 |
| Porechop_ABI, database | 63.2 | 84.2 | 72.7 | 0 | 83.2 | 71.2 | 0 | 0.0 | 0.0 | 0 | 87.0 | 76.9 | 0 | 55.2 | 38.2 | 0 | 98.0 | 96.0 | 0 | 98.4 | 96.9 | 0 | 0.0 | 0.0 | 0 | **0** |
| Porechop_ABI, ab initio | 75.5 | 84.6 | 73.4 | 0 | 83.2 | 71.2 | 0 | 36.8 | 22.8 | 0 | 87.0 | 77.1 | 0 | 55.6 | 38.5 | 0 | 98.2 | 96.4 | 0 | 98.5 | 97.1 | 0 | 60.1 | 43.2 | 0 | 1 |
| fastplong, all | 27.9 | 19.4 | 77.5 | 2,757 | 3.7 | 42.5 | 9,574 | 47.8 | 40.2 | 284 | 6.9 | 56.9 | 6,850 | 65.8 | 79.3 | 600 | 63.4 | 90.7 | 939 | 16.3 | 58.6 | 5,207 | 0.0 | 0.8 | 99,064 | n/a |
| fastplong, outermost | n/a | 47.1 | 86.7 | 803 | 4.8 | 4.4 | 350 | n/a | n/a | n/a | 48.0 | 87.0 | 803 | 52.5 | 41.4 | 157 | 15.3 | 8.4 | 14 | 51.1 | 40.2 | 160 | 89.4 | **94.4** | 71 | n/a |
| fastplong, auto | 16.5 | 22.4 | 77.7 | 2,287 | 0.4 | 2.9 | 5,713 | 0.0 | 0.0 | 955 | 21.9 | 74.4 | 2,315 | 23.0 | 21.2 | 610 | 0.0 | 0.0 | 709 | 45.0 | 83.4 | 1,742 | 19.2 | 71.8 | 2,978 | 2,145 |
| cutadapt, all | 0.0 | 0.0 | 0.0 | 0 | 0.0 | 0.0 | 0 | 0.0 | 0.0 | 0 | 0.0 | 0.0 | 0 | 0.0 | 0.0 | 0 | 0.0 | 0.0 | 0 | 0.0 | 0.0 | 0 | 0.0 | 0.0 | 0 | n/a |
| cutadapt, outermost | 0.0 | 0.0 | 0.0 | 0 | 0.0 | 0.0 | 0 | 0.0 | 0.0 | 0 | 0.0 | 0.0 | 0 | 0.0 | 0.0 | 0 | 0.0 | 0.0 | 0 | 0.0 | 0.0 | 0 | 0.0 | 0.0 | 0 | n/a |
| dorado trim | 0.0 | 0.0 | 0.0 | 0 | 0.0 | 0.0 | 0 | 0.0 | 0.0 | 0 | 0.0 | 0.0 | 0 | 0.0 | 0.0 | 0 | 0.0 | 0.0 | 0 | 0.0 | 0.0 | 0 | 0.0 | 0.0 | 0 | **0** |

*PacBio*

| Tool | Mean F1 (%) | A | | | B | | | P | | | AB | | | AP | | | BP | | | ABP | | | N | | | Control |
|---|---|---|---|---|---|---|---|---|---|---|---|---|---|---|---|---|---|---|---|---|---|---|---|---|---|---|
| | | F1 (%) | Both kept (%) | False splits | F1 (%) | Both kept (%) | False splits | F1 (%) | Both kept (%) | False splits | F1 (%) | Both kept (%) | False splits | F1 (%) | Both kept (%) | False splits | F1 (%) | Both kept (%) | False splits | F1 (%) | Both kept (%) | False splits | F1 (%) | Both kept (%) | False splits | |
| whittle, FASTA | **95.0** | **99.5** | **99.1** | 0 | **71.0** | **55.6** | 4 | 99.0 | 98.1 | 0 | 99.5 | **99.2** | 1 | 99.6 | 99.2 | 0 | 96.1 | 98.0 | 60 | 96.5 | 98.7 | 54 | **98.5** | **97.0** | 0 | n/a |
| whittle, preset | 40.1 | 99.4 | 99.0 | 1 | 0.0 | 0.0 | 0 | 0.0 | 0.0 | 0 | **99.6** | **99.2** | 0 | 61.7 | 44.6 | 0 | 0.0 | 0.0 | 0 | 59.9 | 42.8 | 0 | 0.0 | 0.0 | 1 | 1 |
| whittle, discovery | 91.7 | **99.5** | **99.1** | 0 | 67.8 | 51.5 | 2 | **99.1** | **98.3** | 0 | 99.5 | **99.2** | 1 | 72.2 | 57.3 | 0 | **99.0** | **98.1** | 1 | **100.0** | **100.0** | 0 | 96.4 | 94.1 | 5 | **0** |
| whittle, preset and discovery | 91.1 | 99.4 | 99.0 | 1 | 67.3 | 51.0 | 2 | **99.1** | **98.3** | 0 | 99.5 | 99.1 | 0 | **99.9** | **99.8** | 0 | 79.4 | 70.6 | 1 | 99.9 | 99.8 | 0 | 84.4 | 75.2 | 6 | 1 |
| Porechop_ABI, database | 0.0 | 0.0 | 0.0 | 0 | 0.0 | 0.0 | 0 | 0.0 | 0.0 | 0 | 0.0 | 0.0 | 0 | 0.0 | 0.0 | 0 | 0.0 | 0.0 | 0 | 0.0 | 0.0 | 0 | 0.0 | 0.0 | 0 | **0** |
| Porechop_ABI, ab initio | 41.3 | 53.5 | 36.7 | 3 | 0.7 | 0.3 | 0 | 39.1 | 24.3 | 0 | 76.9 | 62.5 | 0 | 26.5 | 15.3 | 0 | 33.2 | 19.9 | 0 | 26.3 | 15.2 | 0 | 74.4 | 59.3 | 0 | **0** |
| fastplong, all | 13.6 | 6.6 | 58.9 | 7,199 | 0.0 | 0.0 | 178,558 | 54.9 | 68.1 | 774 | 0.0 | 0.0 | 180,185 | 47.6 | 73.6 | 1,305 | 0.0 | 0.0 | 35,138 | 0.0 | 0.0 | 39,207 | 0.0 | 0.9 | 90,928 | n/a |
| fastplong, outermost | n/a | 98.4 | 96.8 | 0 | 0.1 | 2.1 | 15,871 | n/a | n/a | n/a | 98.7 | 97.4 | 0 | 61.4 | 44.3 | 0 | 4.1 | 7.7 | 2,680 | 59.3 | 42.1 | 0 | 85.4 | 95.2 | 124 | n/a |
| fastplong, auto | 12.0 | 6.6 | 58.8 | 7,199 | 0.0 | 0.0 | 0 | 18.5 | 20.3 | 953 | 7.7 | 63.5 | 6,340 | 30.2 | 40.4 | 1,220 | 18.3 | 20.3 | 998 | 0.0 | 0.0 | 0 | 14.6 | 70.5 | 4,131 | 2,285 |
| cutadapt, all | 0.0 | 0.0 | 0.0 | 0 | 0.0 | 0.0 | 0 | 0.0 | 0.0 | 0 | 0.0 | 0.0 | 0 | 0.0 | 0.0 | 0 | 0.0 | 0.0 | 0 | 0.0 | 0.0 | 0 | 0.0 | 0.0 | 0 | n/a |
| cutadapt, outermost | 0.0 | 0.0 | 0.0 | 0 | 0.0 | 0.0 | 0 | 0.0 | 0.0 | 0 | 0.0 | 0.0 | 0 | 0.0 | 0.0 | 0 | 0.0 | 0.0 | 0 | 0.0 | 0.0 | 0 | 0.0 | 0.0 | 0 | n/a |

#### S4.3 Real reads compared with dorado and lima

We scored each tool's trimming of real reads against a reference, the trimming that dorado (ONT) or lima (PacBio) applied to the same reads. Where the cuts disagreed, we searched the bases between them for kit sequence. Table S4 gives the results of every tool configuration on every dataset.

**ONT data.** All four ONT datasets come from the ONT Open Data registry.

- LSK114 (SQK-LSK114): genomic DNA of HG002 from dataset `giab_2025.01`, flow cell PAW70337.
- MAB114 (SQK-MAB114-24): dataset `zymo_16s_2025.09`, reads of the Zymo 16S mock communities MSplus and gutMock from three flow cells.
- RBK114-96 (SQK-RBK114-96): a single rapid barcoding run with 96 barcodes, dataset `nomiss_96BC_P2I_SUP_2026`.
- PCS114 (SQK-PCS114): cDNA of HG002 from dataset `UHRR_HG002_2026.06`.

From each dataset, we selected a subset of the POD5 signal files, adding files in order of increasing size until their estimated yield reached 1 Gb. The subsets gave 1.1 to 1.5 Gb of basecalled sequence each.

dorado 2.1.0 basecalled each subset twice from the same signal, both times with the `sup@v5.2.0` models and `--emit-moves`. The run with `--no-trim` gave the untrimmed reads that every tool received, and the run with default trimming gave the reference. In the barcoded libraries, MAB114 and RBK114-96, `dorado demux` 2.1.0 also removed the barcodes from the reference reads.

Both runs used dorado's default read splitting, so the untrimmed input already held segments of split reads. These segments carry the `pi` tag and make up 504 of the 56,801 records in LSK114, 199 of 697,498 in MAB114, 20,966 of 430,790 in RBK114-96 and 437 of 601,872 in PCS114. The ONT splitting results concern only chimeras that dorado had left whole.

**PacBio data.** Four of the eight PacBio datasets are the Revio libraries with per-base kinetics shown in Figures 1B and 1D. Each contributed about 0.5 Gb (0.35 Gb for Microbial), taken from the start of the published file.

- WGS: movie `m84046_251223_003203_s1` from the Human Pangenome Reference Consortium (Revio SPRQ, December 2025), blood genomic DNA of HG08434 in a SMRTbell prep kit 3.0 library with the barcoded SMRTbell adapter bc2077.
- Fiber-seq: movie `m84400_260514_233654_s3` from the same consortium (Revio SPRQ-Nx, May 2026), a Fiber-seq library of HG02056 with the barcoded SMRTbell adapter bc2003.
- Ampli-Fi: path `revio/2025Q2/microbes/metagenomics` from PacBio (Revio SPRQ, June 2025), the ZymoBIOMICS gut microbiome standard D6331, amplified by PacBio's Ampli-Fi protocol and carrying Twist TruSeq-compatible unique dual index adapters.
- Microbial: run ERR16674849 at the European Nucleotide Archive (Revio SPRQ, September 2025), an isolate of *Streptomyces nigrescens* with the barcoded SMRTbell adapter bc2043.

The four datasets without kinetics, compared in the same way, come from PacBio's download site (https://downloads.pacbcloud.com/public/). Each contributed about 1.3 Gb, again taken from the start of the published file.

- SPRQ-Nx: path `2026Q2/HG002-SPRQ-Nx/Use3`, barcode bc2004 (Revio SPRQ-Nx, May 2026), HG002 genomic DNA with barcoded SMRTbell adapters.
- Vega: path `revio/2025Q2/microbes/isolates-vega` (Vega instrument, June 2025), 16 bacterial isolates barcoded with 96 pairs of Twist TruSeq-compatible adapters. We took the subset in equal shares from each pair's file.
- Kinnex 16S: path `dataset/Kinnex-16S` (Revio SPRQ-Nx, June 2026), 16S amplicons from ZymoBIOMICS and ATCC mock communities, carrying 384 barcoded primer pairs.
- Kinnex RNA: path `dataset/Kinnex-full-length-RNA` (Revio SPRQ, October 2024), full-length Universal Human Reference RNA, carrying barcoded Iso-Seq primers.

The untrimmed reads of Kinnex 16S and Kinnex RNA are segments cut out of Kinnex arrays by skera, and these segments still carry their barcoded primers. The other six datasets were published demultiplexed, and lima-undo 26.2.1 rebuilt their untrimmed reads from the clipped bases that lima records in its tags. In the four libraries with kinetics, lima-undo also restored the kinetic values and modification calls of these bases from the `ls` tag.

We made the PacBio reference by running lima 26.2.1 on the untrimmed reads with the settings that PacBio recommends for each library type. These settings were `--hifi-preset SYMMETRIC-ADAPTERS` for the libraries with barcoded SMRTbell adapters (WGS, Fiber-seq, Microbial and SPRQ-Nx) and `--hifi-preset ASYMMETRIC` for Ampli-Fi, Vega and Kinnex 16S. For Kinnex RNA, we used `--isoseq --peek-guess`.

**Kit sequences.** We searched for the kit sequences of each dataset when scoring read ends and gave them to the "FASTA" configurations of Table S4. For an ONT dataset, the kit sequences are the full contents of whittle's preset for the kit: its adapters, primers, barcodes and flanks. WGS, Fiber-seq, Microbial and SPRQ-Nx share the 96 barcoded adapters (bc2001 to bc2096) of PacBio's SMRTbell adapter plate 3.0, and the Kinnex datasets use PacBio's published primer files. No published file lists the adapters of Ampli-Fi and Vega, so we used the sequences most often clipped by lima, determined separately for each barcode pair in Vega.

When scoring, we also searched for sequences not supplied to any tool: the SMRTbell adapter in all eight PacBio datasets and PacBio's Kinnex array adapters in the two Kinnex datasets. In two datasets, the search further included sequences missing from the kit catalogues. In Kinnex 16S, these were the 16S primers of PacBio's HiFi-16S-workflow, 27F

(AGRGTTYGATYMTGGCTCAG) and 1492R (RGYTACCTTGTTACGACTT). lima leaves both primers in place for the downstream steps of the workflow. In PCS114, these sequences were ONT barcode BC01 and the PCR barcode flank beside it (ATCGCCTACCGTGAC), both of which precede the PCR primer in the reads. An exact copy of BC01 lies in the outermost 200 bases of 73% of PCS114 read ends.

**Reference positions.** We located the reference's cuts by placing each dorado-trimmed read in its untrimmed version with exact search. If a sequence occurs more than once in a read, exact search takes the first copy. We excluded reads without a trimmed record and reads with different basecalls in the two runs. These excluded reads made up 0.3% of the reads across the four ONT datasets and 1.3% of those in LSK114.

We placed lima's records by the same exact search among the untrimmed reads of their own ZMW (zero-mode waveguide). Reads that lima did not assign to a barcode had no reference. These made up 0.3% of the reads in WGS, 0.5% in Fiber-seq, 15.8% in Ampli-Fi, 0.2% in Microbial, 0.3% in SPRQ-Nx, 15.8% in Vega, 0.4% in Kinnex 16S and 2.7% in Kinnex RNA.

We checked our lima runs against the published records, which the instrument had demultiplexed. In WGS, Fiber-seq, Ampli-Fi and Microbial, 183,791 reads were assigned to a barcode by both our runs and the instrument, and 183,790 of these reads were trimmed identically. Our run left 11,806 Ampli-Fi reads unassigned. Given the kit sequences, whittle retained exactly the same bases as the instrument in 11,794 of these reads.

**Tool runs.** All tools received the same untrimmed reads, whittle in uBAM and the others as a FASTQ export. No run applied quality processing or kept segments shorter than 20 bases. Each tool ran once with sequences supplied to it or built into it and once with sequences inferred from the reads, and whittle ran a third time with both. On ONT reads, whittle used the kit preset, discovery or both, always with `--update-moves`. The whittle records for the move-table checks of Section S4.4 came from the kit-preset runs. On PacBio reads, whittle used lima's kit sequences in a FASTA file (the kit FASTA), discovery or both. fastplong ran with the kit sequences as a FASTA file or in automatic mode. Porechop_ABI, which never received the kit sequences, ran with its built-in database or in ab initio mode.

In the kit FASTA, the header descriptions set the role of each sequence, and this role decides when the sequence may split a read (Section S3.1). The barcoded SMRTbell adapters had the barcode role, the Kinnex primers the primer role and every other kit sequence the adapter role.

**Scoring.** We scored each read in four steps:

1. Exact search placed each output segment of the tool in its parent read.
2. We compared the tool's front cut (the start of its first segment) and rear cut (the end of its last segment) with the reference's cuts.
3. For a cut more than three bases from the reference's cut, we searched the bases between the two cuts (the disputed stretch) for kit sequences. The search also included 12 context bases on either side and allowed at most 20% edits.
4. We searched each gap between two segments of a split read in the same way ("Split gaps with kit" in Table S4).

A kit match was a whole kit sequence anywhere in the searched region. In the case of a disputed stretch of 10 or more bases, a kit match could also be a partial match, in which the stretch matched part of one kit sequence. We searched for kit sequences in both orientations, and an IUPAC code in a kit sequence matched any of the bases it stands for. The sequences not supplied to any tool counted as kit sequences in the same way as the supplied ones.

Each end of a read that the tool kept falls into exactly one of five categories, the percentage columns of Table S4:

- "Same": the tool's cut lies within three bases of the reference's.
- "Further, kit": the tool removes bases that the reference kept, and its cut lies no more than three bases past the cover of the kit matches (see below).
- "Kit left": the tool keeps bases that the reference removed, and the search finds a kit match.
- "Further, no kit": the tool removes bases that the reference kept, and either the search finds no kit match or the cut lies more than three bases past the cover.
- Reference further (the "dorado further" or "lima further" column): the tool keeps bases that the reference removed, and the search finds no kit match.

A kit match points to technical sequence somewhere in the searched region, but an end without one does not show that the tool removed sample bases.

The cover is the part of the extra bases that kit matches account for. Because an end counts as "Further, kit" only if the tool's cut lies no more than three bases beyond the cover, a kit match cannot justify trimming into the insert beside it. Starting from the kit match with the fewest edits per base, the cover grows towards the tool's cut:

- The cover extends to the end of another kit sequence that matches as a whole and begins no more than 12 bases beyond the end of the cover.
- The cover extends to the tool's cut when the uncovered bases, either alone or together with the last kit match, match part of a single kit sequence.

For example, consider a 16S read in which the 20-base 27F primer remains at the front after the reference's trimming. When a tool removes only the primer, this end is classified as "Further, kit". If the tool also removes the following 30 gene bases, none of which match a kit sequence, the end is classified as "Further, no kit".

At noisy ONT read ends, the cover can stop between two kit matches if a stretch of more than 12 bases between them is garbled by sequencing errors or absent from the catalogue. Such breaks raise the share of "Further, no kit" ends, and partial matches that arise by chance within 20% edits can lower it.

We classed dropped reads, those without any output segment, by the sequence that the reference kept. A read counted as technical if at least half of the 30-base windows of this sequence (starting every 10 bases) held a kit match, and as insert otherwise. We treated a sequence shorter than 30 bases as a single window. Table S4 omits reads in which the reference kept fewer than 20 bases, as every run's minimum output length excludes them.

A garbled adapter is an adapter with many sequencing errors. If no kit sequence matches a garbled adapter, its removal can give a "Further, no kit" end. The garbled-adapter label marks a "Further, no kit" end at which the 10 removed bases closest to the sample sequence lie within one edit of the sample-facing end of a kit sequence. This label leaves the category of the end unchanged.

**Results.** On LSK114, whittle's preset trimmed further than dorado without a kit match at 238 read ends, and 207 of these ends received the garbled-adapter label. Some of fastplong's trimming beyond the reference comes from its default extension of 10 bases past each adapter. In ab initio mode, Porechop_ABI mistook an Alu element for the rear adapter of LSK114 reads and split 46.7% of the 56,070 reads in the comparison.

dorado assigned a barcode to 5% of the RBK114-96 reads, and Table S4 lists these reads ("Barcode assigned") separately from the rest ("Unclassified"). On the unclassified reads, whittle's preset matched dorado at 95.55% of ends. The preset also removed barcode sequence left in place by dorado, and such ends counted as "Further, kit".

On the 20,215 barcode-assigned reads, the preset agreed with dorado at 52.42% of ends. At 42.70% of their front ends, the preset kept sequence that dorado had removed, with a kit match at 15.62% and without one at 27.07%. At 44.87% of their rear ends, it cut off kit sequence that dorado had kept. The preset dropped 8,746 of these reads, of which 7,318 were classed as technical and 1,428 as insert. On this group, fastplong and Porechop_ABI matched dorado's cuts at 13.10% to 45.37% of read ends. Discovery (Section S3.2) showed that, in this library, the bases preceding the barcode did not match the published flank `GCTTGGGTGTTTAACC`. Instead, discovery identified `AAAACTGGGTGTTTAACC` at this position in 38% of read ends.

In PCS114, Kinnex 16S and Kinnex RNA, some of the sequences that discovery inferred differed from what dorado or lima removed. When whittle received lima's kit sequences instead, it matched lima's trimming at 99.3% to 100% of read ends in each PacBio dataset without kinetics (Table S4).

In PCS114 reads, discovery removed the kit sequences and the UMI but, unlike dorado, mostly kept the four T bases after the PCR primer.

Discovery recovered parts of both 16S primers, 27F and 1492R, from Kinnex 16S reads, finding each at 86% to 87% of the sampled read ends. None of the kit's 44 barcodes (12 forward and 32 reverse, combined into its 384 pairs) was discovered. However, at 91.8% of read ends, trimming at the primers also removed the barcodes. The discovered 1492R sequence (`AAGTCGTAACAAGGTARC`) differs by three edits from `AAGTCGAACGGTAAC`, a conserved 16S motif. At 5.3% of ends, discovery trimmed into the gene at this motif, on average 66 bases beyond lima's cut, that is, 46 bases past the 27F primer.

In Kinnex RNA reads, discovery inferred a 3′ sequence in which 15 adenines precede the Iso-Seq 3′ primer. Because lima keeps the poly(A) tail, discovery trimmed beyond lima at 50.6% of ends, by 25 bases on average and without a kit match. The deeper layers inferred by discovery were fragments of Alu elements, each identified at 1% to 4% of the sampled read ends. Splitting at these fragments divided 6.6% of the reads and left 59,752 gaps between segments, of which 3,244 contained a kit match.

**Table S4.** Trimming of real reads relative to dorado (ONT) and lima (PacBio). The "Reads" column counts the parent reads compared, including dropped reads, which have no output segment. Percentages give the share of read ends in each category of Section S4.3, pooling both ends of every parent read that the tool kept. The columns "Dropped, technical" and "Dropped, insert" count the dropped reads in each class. The column "Split gaps with kit" gives x of y, where y is the number of gaps between split segments and x is the number of these gaps that contain a kit match. The "FASTA" runs received the kit sequences in a FASTA file, and the "whittle both" runs combined discovery with the preset (ONT) or the kit FASTA (PacBio). The other configuration labels ("preset", "discovery", "auto" for automatic mode, "database" and "ab initio") are those of Table S2. The second ONT group divides the RBK114-96 reads by dorado's barcode assignment ("Barcode assigned" and "Unclassified"). The PacBio groups list the four libraries of Figure 1D, which carry kinetics, above the four datasets without kinetics.

*ONT, relative to dorado*

| Kit | Tool | Reads | Same (%) | Further, kit (%) | Kit left (%) | Further, no kit (%) | dorado further (%) | Split gaps with kit | Dropped, technical | Dropped, insert |
|---|---|---|---|---|---|---|---|---|---|---|
| LSK114 | whittle preset | 56,070 | 91.95 | 7.71 | 0.06 | 0.21 | 0.06 | 54 of 54 | 1 | 0 |
| | whittle discovery | 56,070 | 93.39 | 5.97 | 0.41 | 0.21 | 0.02 | 50 of 50 | 1 | 0 |
| | whittle both | 56,070 | 91.92 | 7.76 | 0.05 | 0.21 | 0.07 | 54 of 54 | 1 | 0 |
| | fastplong FASTA | 56,070 | 58.95 | 2.18 | 0.17 | 38.70 | 0.00 | 1,354 of 41,703 | 1 | 1 |
| | fastplong auto | 56,070 | 60.31 | 2.19 | 0.33 | 37.17 | 0.00 | 231 of 801 | 1 | 0 |
| | Porechop_ABI database | 56,070 | 50.30 | 16.77 | 0.40 | 32.53 | 0.00 | 27 of 27 | 1 | 8 |
| | Porechop_ABI ab initio | 56,070 | 36.33 | 16.58 | 0.18 | 46.91 | 0.00 | 80 of 66,863 | 1 | 74 |
| MAB114 | whittle preset | 696,241 | 49.62 | 49.26 | 0.33 | 0.41 | 0.39 | 14,662 of 14,662 | 6 | 7 |
| | whittle discovery | 696,241 | 47.66 | 46.78 | 2.66 | 2.33 | 0.57 | 11,446 of 11,475 | 2 | 6 |

| Kit | Tool | Reads | Same (%) | Further, kit (%) | Kit left (%) | Further, no kit (%) | dorado further (%) | Split gaps with kit | Dropped, technical | Dropped, insert |
|---|---|---|---|---|---|---|---|---|---|---|
| | whittle both | 696,241 | 49.45 | 49.43 | 0.16 | 0.41 | 0.54 | 14,304 of 14,304 | 6 | 9 |
| | fastplong FASTA | 696,241 | 25.71 | 43.21 | 0.11 | 30.66 | 0.31 | 16,193 of 48,216 | 47 | 250 |
| | fastplong auto | 696,241 | 30.17 | 43.28 | 1.66 | 24.71 | 0.18 | 8,449 of 21,494 | 2 | 72 |
| | Porechop_ABI database | 696,241 | 55.06 | 0.15 | 44.28 | 0.18 | 0.34 | 1 of 1 | 1 | 57 |
| | Porechop_ABI ab initio | 696,241 | 31.54 | 8.35 | 33.92 | 25.86 | 0.33 | 2,705 of 2,782 | 19 | 150 |
| RBK114-96 | whittle preset | 423,260 | 94.30 | 2.97 | 0.84 | 0.32 | 1.56 | 4,268 of 4,268 | 17,950 | 6,974 |
| | whittle discovery | 423,260 | 93.90 | 2.40 | 0.74 | 1.48 | 1.49 | 1,612 of 1,792 | 18,281 | 12,427 |
| | whittle both | 423,260 | 94.62 | 2.30 | 0.58 | 1.00 | 1.50 | 1,999 of 2,205 | 19,413 | 8,816 |
| | fastplong FASTA | 423,260 | 48.22 | 33.43 | 0.14 | 18.00 | 0.21 | 8,800 of 138,212 | 19,500 | 7,932 |
| | fastplong auto | 423,260 | 51.11 | 35.28 | 0.42 | 13.12 | 0.07 | 1,708 of 2,303 | 17,183 | 5,830 |
| | Porechop_ABI database | 423,260 | 48.17 | 12.00 | 0.88 | 38.74 | 0.20 | 1,469 of 1,476 | 8,530 | 9,379 |
| | Porechop_ABI ab initio | 423,260 | 48.53 | 11.88 | 0.57 | 38.79 | 0.22 | 901 of 902 | 18,212 | 17,847 |
| PCS114 | whittle preset | 600,347 | 90.73 | 5.72 | 0.10 | 1.18 | 2.27 | 25,078 of 25,078 | 16 | 44 |
| | whittle discovery | 600,347 | 48.46 | 4.76 | 0.77 | 1.90 | 44.10 | 22,422 of 22,482 | 44 | 151 |
| | whittle both | 600,347 | 90.14 | 5.75 | 0.04 | 1.80 | 2.26 | 21,585 of 21,625 | 119 | 241 |
| | fastplong FASTA | 600,347 | 4.69 | 5.20 | 42.91 | 24.45 | 22.74 | 26,478 of 136,752 | 70 | 183 |
| | fastplong auto | 600,347 | 5.51 | 4.81 | 89.28 | 0.35 | 0.06 | 22,363 of 23,748 | 25 | 67 |
| | Porechop_ABI database | 600,347 | 4.87 | 5.56 | 41.50 | 1.57 | 46.51 | 16,061 of 16,066 | 153 | 726 |
| | Porechop_ABI ab initio | 600,347 | 4.78 | 5.56 | 41.50 | 1.65 | 46.51 | 16,049 of 16,054 | 157 | 730 |

*ONT RBK114-96 by dorado barcode assignment, relative to dorado*

| Subset | Tool | Reads | Same (%) | Further, kit (%) | Kit left (%) | Further, no kit (%) | dorado further (%) | Split gaps with kit | Dropped, technical | Dropped, insert |
|---|---|---|---|---|---|---|---|---|---|---|
| Barcode assigned | whittle preset | 20,215 | 52.42 | 24.74 | 8.02 | 0.38 | 14.44 | 1,151 of 1,151 | 7,318 | 1,428 |
| | whittle discovery | 20,215 | 54.22 | 19.75 | 7.98 | 4.97 | 13.09 | 206 of 243 | 7,691 | 2,464 |
| | whittle both | 20,215 | 55.65 | 17.66 | 6.31 | 5.16 | 15.22 | 146 of 192 | 8,300 | 2,511 |
| | fastplong FASTA | 20,215 | 43.94 | 30.84 | 1.97 | 14.46 | 8.80 | 111 of 2,082 | 8,631 | 2,001 |
| | fastplong auto | 20,215 | 45.37 | 32.95 | 9.18 | 10.10 | 2.39 | 523 of 820 | 6,751 | 1,163 |
| | Porechop_ABI database | 20,215 | 14.17 | 32.91 | 12.35 | 34.74 | 5.84 | 15 of 15 | 3,811 | 3,341 |
| | Porechop_ABI ab initio | 20,215 | 13.10 | 34.08 | 8.50 | 34.44 | 9.88 | 6 of 7 | 8,531 | 4,152 |
| Unclassified | whittle preset | 403,045 | 95.55 | 2.33 | 0.63 | 0.32 | 1.18 | 3,117 of 3,117 | 10,632 | 5,546 |
| | whittle discovery | 403,045 | 94.94 | 1.94 | 0.55 | 1.38 | 1.18 | 1,406 of 1,549 | 10,590 | 9,963 |
| | whittle both | 403,045 | 95.57 | 1.93 | 0.44 | 0.90 | 1.16 | 1,853 of 2,013 | 11,113 | 6,305 |
| | fastplong FASTA | 403,045 | 48.33 | 33.50 | 0.09 | 18.09 | 0.00 | 8,689 of 136,130 | 10,869 | 5,931 |
| | fastplong auto | 403,045 | 51.29 | 35.36 | 0.14 | 13.21 | 0.00 | 1,185 of 1,483 | 10,432 | 4,667 |
| | Porechop_ABI database | 403,045 | 49.31 | 11.31 | 0.50 | 38.87 | 0.01 | 1,454 of 1,461 | 4,719 | 6,038 |
| | Porechop_ABI ab initio | 403,045 | 49.23 | 11.44 | 0.42 | 38.88 | 0.03 | 895 of 895 | 9,681 | 13,695 |

*PacBio, relative to lima*

| Library | Tool | Reads | Same (%) | Further, kit (%) | Kit left (%) | Further, no kit (%) | lima further (%) | Split gaps with kit | Dropped, technical | Dropped, insert |
|---|---|---|---|---|---|---|---|---|---|---|
| WGS | whittle FASTA | 22,590 | 99.20 | 0.33 | 0.25 | 0.00 | 0.23 | 7 of 7 | 0 | 0 |
| | whittle discovery | 22,590 | 99.47 | 0.13 | 0.17 | 0.01 | 0.23 | 3 of 3 | 0 | 0 |
| | whittle both | 22,590 | 99.20 | 0.33 | 0.25 | 0.00 | 0.23 | 7 of 7 | 0 | 0 |
| | fastplong FASTA | 22,590 | 2.11 | 11.01 | 21.13 | 65.66 | 0.09 | 340,261 of 935,985 | 0 | 0 |
| | fastplong auto | 22,590 | 4.99 | 0.00 | 94.79 | 0.00 | 0.22 | 0 of 0 | 0 | 0 |
| | Porechop_ABI database | 22,590 | 4.99 | 0.00 | 94.79 | 0.00 | 0.22 | 0 of 0 | 0 | 0 |
| | Porechop_ABI ab initio | 22,590 | 17.40 | 6.07 | 3.63 | 72.75 | 0.15 | 0 of 0 | 0 | 0 |
| Fiber-seq | whittle FASTA | 21,444 | 99.20 | 0.33 | 0.21 | 0.00 | 0.26 | 10 of 10 | 0 | 0 |
| | whittle discovery | 21,444 | 99.34 | 0.26 | 0.14 | 0.00 | 0.26 | 1 of 1 | 0 | 0 |
| | whittle both | 21,444 | 99.20 | 0.33 | 0.21 | 0.00 | 0.26 | 10 of 10 | 0 | 0 |
| | fastplong FASTA | 21,444 | 0.65 | 12.06 | 10.90 | 76.27 | 0.12 | 341,526 of 940,594 | 0 | 0 |
| | fastplong auto | 21,444 | 1.58 | 0.00 | 98.16 | 0.00 | 0.26 | 0 of 0 | 0 | 0 |
| | Porechop_ABI database | 21,444 | 1.58 | 0.00 | 98.16 | 0.00 | 0.26 | 0 of 0 | 0 | 0 |
| | Porechop_ABI ab initio | 21,444 | 11.84 | 7.97 | 1.82 | 78.27 | 0.11 | 0 of 0 | 0 | 0 |
| Ampli-Fi | whittle FASTA | 62,787 | 100.00 | 0.00 | 0.00 | 0.00 | 0.00 | 15 of 15 | 0 | 0 |
| | whittle discovery | 62,787 | 99.41 | 0.02 | 0.00 | 0.56 | 0.01 | 15 of 15 | 0 | 0 |
| | whittle both | 62,787 | 100.00 | 0.00 | 0.00 | 0.00 | 0.00 | 15 of 15 | 0 | 0 |
| | fastplong FASTA | 62,787 | 0.00 | 12.99 | 50.00 | 37.01 | 0.00 | 14 of 15 | 0 | 0 |
| | fastplong auto | 62,787 | 0.00 | 0.00 | 100.00 | 0.00 | 0.00 | 0 of 0 | 0 | 0 |
| | Porechop_ABI database | 62,787 | 0.00 | 0.00 | 100.00 | 0.00 | 0.00 | 0 of 0 | 0 | 0 |
| | Porechop_ABI ab initio | 62,787 | 29.99 | 0.18 | 0.00 | 69.83 | 0.00 | 16 of 16 | 0 | 0 |
| Microbial | whittle FASTA | 76,970 | 99.20 | 0.23 | 0.29 | 0.00 | 0.29 | 3 of 3 | 0 | 0 |
| | whittle discovery | 76,970 | 99.19 | 0.28 | 0.23 | 0.01 | 0.29 | 3 of 3 | 0 | 0 |

| Library | Tool | Reads | Same (%) | Further, kit (%) | Kit left (%) | Further, no kit (%) | lima further (%) | Split gaps with kit | Dropped, technical | Dropped, insert |
|---|---|---|---|---|---|---|---|---|---|---|
| | whittle both | 76,970 | 99.20 | 0.23 | 0.29 | 0.00 | 0.29 | 3 of 3 | 0 | 0 |
| | fastplong FASTA | 76,970 | 2.34 | 15.10 | 29.50 | 52.89 | 0.17 | 109,867 of 369,009 | 0 | 0 |
| | fastplong auto | 76,970 | 3.71 | 0.00 | 96.00 | 0.00 | 0.29 | 0 of 0 | 0 | 0 |
| | Porechop_ABI database | 76,970 | 3.71 | 0.00 | 96.00 | 0.00 | 0.29 | 0 of 0 | 0 | 0 |
| | Porechop_ABI ab initio | 76,970 | 6.87 | 6.65 | 1.61 | 84.71 | 0.16 | 1 of 1 | 0 | 0 |

*PacBio, datasets without kinetics, relative to lima*

| Library | Tool | Reads | Same (%) | Further, kit (%) | Kit left (%) | Further, no kit (%) | lima further (%) | Split gaps with kit | Dropped, technical | Dropped, insert |
|---|---|---|---|---|---|---|---|---|---|---|
| SPRQ-Nx | whittle FASTA | 66,353 | 99.27 | 0.34 | 0.20 | 0.00 | 0.20 | 41 of 41 | 0 | 0 |
| | whittle discovery | 66,353 | 99.05 | 0.63 | 0.12 | 0.00 | 0.20 | 1 of 1 | 0 | 0 |
| | whittle both | 66,353 | 99.27 | 0.34 | 0.20 | 0.00 | 0.20 | 41 of 41 | 0 | 0 |
| | fastplong FASTA | 66,353 | 0.62 | 11.71 | 10.89 | 76.70 | 0.08 | 878,259 of 2,422,821 | 0 | 0 |
| | fastplong auto | 66,353 | 1.46 | 0.00 | 98.34 | 0.00 | 0.20 | 0 of 0 | 0 | 0 |
| | Porechop_ABI database | 66,353 | 1.46 | 0.00 | 98.34 | 0.00 | 0.20 | 0 of 0 | 0 | 0 |
| | Porechop_ABI ab initio | 66,353 | 10.34 | 7.55 | 3.45 | 78.51 | 0.15 | 0 of 0 | 0 | 0 |
| Vega | whittle FASTA | 129,105 | 99.99 | 0.00 | 0.00 | 0.00 | 0.01 | 1 of 1 | 0 | 0 |
| | whittle discovery | 129,105 | 99.97 | 0.00 | 0.02 | 0.00 | 0.01 | 0 of 0 | 0 | 0 |
| | whittle both | 129,105 | 99.99 | 0.00 | 0.00 | 0.00 | 0.01 | 1 of 1 | 0 | 0 |
| | fastplong FASTA | 129,105 | 0.00 | 43.49 | 50.00 | 6.50 | 0.00 | 0 of 0 | 0 | 0 |
| | fastplong auto | 129,105 | 0.00 | 0.00 | 100.00 | 0.00 | 0.00 | 0 of 0 | 0 | 0 |
| | Porechop_ABI database | 129,105 | 0.00 | 0.00 | 100.00 | 0.00 | 0.00 | 0 of 0 | 0 | 0 |
| | Porechop_ABI ab initio | 129,105 | 43.16 | 0.37 | 0.03 | 56.43 | 0.00 | 1 of 1 | 0 | 0 |
| Kinnex 16S | whittle FASTA | 830,346 | 99.31 | 0.66 | 0.00 | 0.02 | 0.00 | 49 of 49 | 0 | 0 |
| | whittle discovery | 830,346 | 0.00 | 91.78 | 2.94 | 5.29 | 0.00 | 8 of 8 | 0 | 0 |
| | whittle both | 830,346 | 2.92 | 91.79 | 0.00 | 5.29 | 0.00 | 45 of 45 | 0 | 0 |
| | fastplong FASTA | 830,346 | 17.87 | 32.11 | 32.13 | 17.89 | 0.00 | 44 of 2,255 | 0 | 0 |
| | fastplong auto | 830,346 | 17.87 | 1.04 | 63.20 | 17.89 | 0.00 | 27 of 2,236 | 0 | 0 |
| | Porechop_ABI database | 830,346 | 0.00 | 0.00 | 100.00 | 0.00 | 0.00 | 0 of 0 | 0 | 0 |
| | Porechop_ABI ab initio | 830,346 | 0.00 | 0.00 | 18.20 | 81.79 | 0.00 | 21 of 21 | 0 | 0 |
| Kinnex RNA | whittle FASTA | 692,954 | 99.98 | 0.01 | 0.00 | 0.00 | 0.01 | 20 of 20 | 0 | 0 |
| | whittle discovery | 692,954 | 49.38 | 0.03 | 0.00 | 50.59 | 0.01 | 3,244 of 59,752 | 0 | 3 |
| | whittle both | 692,954 | 99.98 | 0.01 | 0.00 | 0.00 | 0.01 | 20 of 20 | 0 | 0 |
| | fastplong FASTA | 692,954 | 0.00 | 6.05 | 15.22 | 78.73 | 0.00 | 248 of 5,180 | 0 | 0 |
| | fastplong auto | 692,954 | 0.00 | 6.05 | 30.23 | 63.72 | 0.00 | 148 of 4,808 | 0 | 0 |
| | Porechop_ABI database | 692,954 | 0.00 | 0.00 | 100.00 | 0.00 | 0.00 | 0 of 0 | 0 | 0 |
| | Porechop_ABI ab initio | 692,954 | 21.80 | 0.01 | 50.00 | 28.20 | 0.00 | 27 of 27 | 0 | 0 |

### S4.4 Annotation checks

From the untrimmed input, standalone scripts calculated the annotations that each output record should carry. We compared these expected values with whittle's records and with those that dorado, lima or chopper wrote for the same reads. Table S5 covers the real reads of Section S4.3, while Table S6 covers only modification calls, checked on the inputs of the cropping and filtering benchmark (Section S4.5).

**ONT signal tags.** We tested the move table (`mv`) and signal start (`ts`) of whittle's kit-preset records and dorado's reference records for the ONT reads of Section S4.3. A read was included in the check only if whittle wrote a single output segment for it and this segment held exactly the bases retained by dorado. Some of these segments were split records: whittle had split the read and kept only this segment.

All the move tables examined were 0/1 tables (Section S2). The expected `mv` of a segment starts at the entry that marks the segment's first base. It ends just before the entry that marks the first removed base after the segment (Figure S2C). If no base after the segment is removed, the expected `mv` runs to the end of the input table. The expected `ts` is input `ts` + stride × number of entries before this slice. In split records, `sp` + `ts` had to equal input `sp` (0 if absent) + expected `ts`. This condition held in all 712 split records.

The sequence retained in a record can occur more than once in its input read, and each occurrence implies its own expected signal start. The signal start of a record therefore shows which copy the record kept. A whittle record passed if its signal start and move table matched the values expected at any occurrence. In contrast, we tested dorado's records only at the first copy, where exact search had placed them (Section S4.3).

Across the 901,627 records checked in the four libraries, every whittle record passed (Table S5, Figure 1B), and dorado's records matched in 98.3% to 99.9% of reads per library. Table S5 counts a dorado record as differing if it retained a copy other than the first, so dorado's match rates are lower bounds. whittle's records matched a copy other than the first in 5 RBK114-96 and 35 PCS114 reads, under 0.01% of each library.

Table S5 omits the signal endpoint `ns`, which the two tools set by different rules. In a trimmed dorado record, `ns` marks the end of the last move-table block, at `ts` + stride × n for a move table of n entries. whittle sets `ns` to the end of the signal of the kept bases (Section S2). If trimming removes only the start of a DNA read, whittle keeps the exact input value of `ns`.

Among the records of Table S5, the two tools wrote the same `ns` in 96.0% of LSK114, 42.0% of MAB114, 21.7% of RBK114-96 and 96.7% of PCS114 records. In the 712 split records, whittle's `ns` counts only the samples of the record's own signal. In every other record with unequal values, dorado's `ns` ended at its last move-table block. In all of these records, whittle had set `ns` according to its own rule. They include 22 records (5 RBK114-96 and 17 PCS114) reported by the check script as departures from this rule.

These 22 reports are an artefact of how the check script placed the kept sequence. Each of these reads ended in a run of T bases, and both tools kept only the last 23 to 261 bases of this run. Exact search placed this sequence at its first copy, which begins at the start of the run, two bases (RBK114-96) or four bases (PCS114) before the copy at the read end. The script therefore classified each record as a crop that removed bases from the end of the read. For such a crop, the script expected whittle's `ns` to mark the end of the last retained move-table block. However, the signal starts and move tables of both tools matched only the copy at the read end, which shows that both tools had trimmed only the start of the read. whittle's records accordingly carried the input `ns`, 1 to 5 samples above dorado's. All 22 records are among the 40 reads in which whittle's record matched a copy other than the first.

**Table S5.** Annotation checks on reads trimmed to identical bases by whittle and by the reference tool, dorado for ONT and lima for PacBio. Library names follow Section S4.3. For move tables, a record matches if `mv` and the signal start equal the values calculated from the untrimmed input. For a split record, the signal start is its offset in the parent signal, `sp` + `ts`. For kinetics, a record matches if `fi`, `fp`, `ri` and `rp` each equal their slice of the input arrays. For modification calls, a record matches if its calls are those of the input at the retained bases. Both the record's calls and the input calls are decoded with HTSlib (Section S4.4). Values and Calls give the numbers of kinetic values and calls compared.

*ONT, move tables and signal starts*

| Library | Reads checked | whittle matches | whittle differs | dorado matches |
|---|---|---|---|---|
| LSK114 | 11,599 | 11,599 | 0 | 11,402 |
| MAB114 | 66,247 | 66,247 | 0 | 66,152 |
| RBK114-96 | 343,439 | 343,439 | 0 | 342,550 |
| PCS114 | 480,342 | 480,342 | 0 | 478,382 |

*PacBio, kinetics*

| Library | Records checked | Values | whittle matches | whittle differs | lima matches |
|---|---|---|---|---|---|
| WGS | 22,026 | 1,905,034,572 | 22,026 | 0 | 22,026 |
| Fiber-seq | 20,935 | 1,903,895,688 | 20,935 | 0 | 20,935 |
| Ampli-Fi | 62,740 | 1,675,414,930 | 62,740 | 0 | 62,740 |
| Microbial | 75,419 | 1,368,214,172 | 75,419 | 0 | 75,419 |

*PacBio, modification calls*

| Library | Records checked | Calls | whittle matches | whittle differs | lima matches |
|---|---|---|---|---|---|
| WGS | 22,026 | 6,631,202 | 22,026 | 0 | 22,026 |
| Fiber-seq | 20,935 | 46,475,772 | 20,935 | 0 | 20,935 |
| Ampli-Fi | 62,740 | 23,139,087 | 62,740 | 0 | 62,740 |
| Microbial | 75,419 | 47,754,376 | 75,419 | 0 | 75,419 |
| SPRQ-Nx | 64,661 | 14,990,068 | 64,661 | 0 | 64,661 |
| Vega | 129,049 | 66,104,081 | 129,049 | 0 | 129,049 |
| Kinnex 16S | 338,348 | 35,039,538 | 338,348 | 0 | 338,348 |
| Kinnex RNA | 690,813 | 33,555,533 | 690,813 | 0 | 484,497 |

**Modification calls.** HTSlib (Bonfield et al. 2021), called through pysam 0.24.1, decoded the input and output calls alike, independently of whittle's code. For Table S6, four separate, untimed runs processed the first 20,000 records from each input of the cropping and filtering benchmark (Section S4.5):

- whittle with fixed cropping alone, writing uBAM.
- whittle with adapter and quality processing, writing uBAM.
- whittle with the same processing, writing tagged FASTQ.
- chopper with `--update-mods`, cropping a tagged FASTQ export of the records.

All four runs cropped 10 bases from each end of every segment and kept output segments of at least 100 bases. For adapter processing, whittle used the kit preset of each dataset: `lsk114` for ligation reads, `ulk114` for ultra-long reads and `pacbio` for both Revio inputs. whittle's quality processing split segments at every run of 20 or more bases below Q10 (quality method `runs`, Section S1).

The script tested each output record in four steps:

1. Each ONT record was paired with its input read through the parent read name. PacBio records were paired through the movie name and ZMW number instead, because their names can carry query coordinates updated by whittle (Section S2).

2. Exact search placed the record's sequence in the input read at position s, the start of the kept interval.
3. The record's decoded calls had to be the input calls on the retained bases. A call at input position p had to appear at output position p − s, with its canonical base, strand, modification code and probability unchanged. Calls absent from the record, or present only in the record, were mismatches.
4. The record's modification groups, including empty ones, had to be those of the input read. Each group also had to retain its mode, which is marked by the optional `.` (implicit) or `?` (explicit) after its modification codes. Any `MN` value had to equal the record's length.

Exact search found the sequence of every output record in its input read, for some records at two or more positions. We excluded such records, because a call position, unlike a signal start, does not show which copy a record kept. Four such records came from cropping and 2,641 from adapter and quality processing (2,634 ONT ultra-long and 7 ONT ligation records).

In the 232,504 uBAM records from adapter and quality processing, all 68,144,389 kept calls matched their expected values, and the tagged FASTQ output carried the same calls. Cropping alone left 69,139,171 calls in whittle's 78,127 checked records, and every call was as expected. chopper's cropped records reached the same totals without a mismatch. Every record carried the modification groups of its input read, and every `MN` value tested was correct (Table S6).

A looser form of this test compared whittle, given the kit FASTA, with lima on the eight PacBio datasets of Section S4.3. This test used the reads that met two conditions: each tool wrote a single record for the read, and the two records retained identical bases. The calls had to meet the rule of step 3, but `MN` and the implicit or explicit mode of the groups went untested. If trimming removed every call of a group, the group passed whether the tool kept it or removed it.

None of whittle's 1,403,991 records, which held 273,689,657 calls, had a mismatch. lima's records matched the input in seven of the eight datasets. In Kinnex RNA, 206,316 of the 690,813 checked lima records (29.9%; Table S5) carried calls that differed from the input. In its `--isoseq` mode, lima flagged Kinnex RNA records as reverse-strand but left their sequence and modification tags in the original orientation. For a record flagged in this way, HTSlib reads the calls on the opposite strand.

**Kinetic arrays.** For an input read of length L and a kept interval [s,e), the arrays `fi` and `fp` of an output record had to equal the slice [s,e) of the input arrays. The arrays `ri` and `rp` are stored last base first (Section S2, Figure S2D), so they had to equal the slice [L−e,L−s). An empty input array had to stay empty (Section S2).

Of the cropping and filtering inputs, only Revio Fiber-seq carried kinetics, with all four arrays. In the 20,000 uBAM records from cropping alone and the 20,558 records from adapter and quality processing, no array differed from its expected slice. Every nonempty input array was also as long as its read. We requested only `MM`, `ML` and `MN` in the tagged FASTQ outputs, so the kinetic check left these outputs out.

On the four Revio libraries of Section S4.3, whittle (given the kit FASTA) and lima kept identical bases in 181,120 records. In these records, every array of both tools equalled its input slice, across 6,852,559,362 values in all (Table S5).

**Limitations.** The checks did not test move tables after quality processing, in RNA reads or in reads that keep two or more segments. Nor did they cover the read names and query coordinates (`qs`, `qe`) of PacBio records, the poly(A) estimates (`pt`, `pa`), the run-length coverage (`sa`) or the recalculated `du` and `st`.

**Table S6.** Modification-call checks on the four outputs made from each input of the cropping and filtering benchmark (Section S4.5). Output names the tool, the processing (crop for fixed cropping, split for adapter and quality processing) and the format (BAM for uBAM, FASTQ for tagged FASTQ). Checked records counts the records with a sequence located at a single position of the input read. Ambiguous counts the records excluded because their sequence was located at multiple positions. Calls is the number of observed calls, equal in every row to the number expected. Call mismatches counts the records in which a call is missing or extra or differs in probability, code, strand, base or position. MN checked counts the records that carry `MN`, and the `MN` value of each of these records was tested against the output length. MN mismatches counts the records that failed this test.

| Dataset | Output | Checked records | Ambiguous | Calls | Call mismatches | MN checked | MN mismatches |
|---|---|---|---|---|---|---|---|
| ONT ligation | whittle crop, BAM | 18,127 | 4 | 1,604,558 | 0 | 18,127 | 0 |
| ONT ligation | whittle split, BAM | 21,795 | 7 | 1,594,368 | 0 | 21,795 | 0 |
| ONT ligation | whittle split, FASTQ | 21,795 | 7 | 1,594,368 | 0 | 21,795 | 0 |
| ONT ligation | chopper crop, FASTQ | 18,127 | 4 | 1,604,558 | 0 | 18,127 | 0 |
| ONT ultra-long | whittle crop, BAM | 20,000 | 0 | 58,482,800 | 0 | 20,000 | 0 |
| ONT ultra-long | whittle split, BAM | 169,691 | 2,634 | 57,498,840 | 0 | 169,691 | 0 |
| ONT ultra-long | whittle split, FASTQ | 169,691 | 2,634 | 57,498,840 | 0 | 169,691 | 0 |
| ONT ultra-long | chopper crop, FASTQ | 20,000 | 0 | 58,482,800 | 0 | 20,000 | 0 |
| Revio Fiber-seq | whittle crop, BAM | 20,000 | 0 | 4,348,568 | 0 | 20,000 | 0 |
| Revio Fiber-seq | whittle split, BAM | 20,558 | 0 | 4,348,234 | 0 | 20,558 | 0 |
| Revio Fiber-seq | whittle split, FASTQ | 20,558 | 0 | 4,348,234 | 0 | 20,558 | 0 |
| Revio Fiber-seq | chopper crop, FASTQ | 20,000 | 0 | 4,348,568 | 0 | 0 | 0 |
| Revio SPRQ-Nx | whittle crop, BAM | 20,000 | 0 | 4,703,245 | 0 | 19,970 | 0 |
| Revio SPRQ-Nx | whittle split, BAM | 20,460 | 0 | 4,702,947 | 0 | 20,428 | 0 |
| Revio SPRQ-Nx | whittle split, FASTQ | 20,460 | 0 | 4,702,947 | 0 | 20,428 | 0 |
| Revio SPRQ-Nx | chopper crop, FASTQ | 20,000 | 0 | 4,703,245 | 0 | 0 | 0 |

#### S4.5 Cropping and filtering

The cropping and filtering benchmark timed only the steps shared by whittle, chopper and fastplong: a fixed crop at both ends of each read and filters on read length and mean quality. GNU time recorded the wall time, CPU time and peak memory of every run. Table S2 gives the time and memory of adapter trimming and discovery, measured on simulated reads only.

**Inputs.** The inputs were four public uBAM files. The two ONT files contain R10.4.1 reads of HG002, ligation reads in one file and ultra-long reads in the other. Both files come from the data submitted by UCSC and GIAB to the Human Pangenome Reference Consortium (HPRC) under `5b73fa0e-658a-4248-b2b8-cd16155bc157--UCSC_GIAB_R1041_nanopore`. Of the two PacBio files, one contains Revio Fiber-seq reads of HG02624 from a single SMRT Cell in the HPRC data. The other contains HG002 reads from PacBio's Revio SPRQ-Nx Use 3 release (https://downloads.pacbcloud.com/public/2026Q2/HG002-SPRQ-Nx/Use3/hifi_reads/). Of the four files, only the SPRQ-Nx file also supplied a dataset for Section S4.3. The Fiber-seq library in that section comes from a different sample, HG02056.

We discarded ultra-long reads shorter than 70 kb before samtools (seed 11) subsampled the inputs, with all tags kept, to a target of 30 Gb of sequence per file. The Fiber-seq file held less than the target, and we used it whole. The prepared inputs were:

- ONT ligation: `09_09_24_R1041_LIG_HG002_15kb_C1_dorado1.3.1_sup5.2.0_5mCG_5hmCG.bam`, 8,030,530 reads with 29.980 Gb.
- ONT ultra-long: `09_16_24_R1041_UL_HG002_7C_dorado0.9.1_sup5.0.0_5mCG_5hmCG.bam`, 216,065 reads with 30.001 Gb.
- Revio Fiber-seq: `m84081_240501_202516_s2.hifi_reads.bc2089.bam`, 745,544 reads with 16.941 Gb.
- Revio SPRQ-Nx: `m84031_260507_084733_s1.hifi_reads.bc2004.bam`, 1,676,535 reads with 30.015 Gb.

All four inputs carried modification tags, and the Fiber-seq reads also held the kinetic arrays `fi`, `fp`, `ri` and `rp`. Move tables were absent from both ONT inputs, so the move-table checks of Section S4.4 used the ONT datasets of Section S4.3.

**Tool settings.** From each uBAM, we exported four FASTQ files: plain and gzip-compressed (pigz level 6), each with and without the header tags `MM`, `ML` and `MN`. All three tools processed the untagged files. whittle and `chopper --update-mods` processed the tagged files, and only whittle processed the uBAM. Adapter search, quality trimming and splitting were switched off in every run. Each run cropped 10 bases from each end of every read and retained reads of at least 500 bases with a mean quality of at least Q10. For fastplong, `-u 100` disabled the qualified-base fraction filter. The full command lines are in the archive.

Two differences between the tools can change which reads pass the filters. First, whittle (Section S1) and chopper average the error probabilities of the bases in a read and express this mean as a Phred value, while fastplong (`-m 10`) averages the Phred scores themselves. For example, if half the bases of a read are Q10 and half are Q20, the average Phred score is Q15, but the mean error probability is 0.055, equivalent to Q12.6. Second, chopper applies the mean quality filter before cropping, and whittle applies it after cropping. The mean quality calculated by chopper therefore still includes the 10 bases removed from each end by cropping.

Both whittle and fastplong used gzip level 4, and whittle kept its default level of 6 for uBAM output. For gzip FASTQ, we piped chopper's output into `pigz -4 -p T`, where T is the thread count requested from chopper. The compression threads of pigz therefore ran alongside those of chopper.

**Timing.** Every combination of four datasets, four thread counts (1, 4, 8 and 16) and eleven pairings of tool and format ran twice, and all 352 runs completed. For each dataset, format and thread count, the tools ran back to back in a fixed order, whittle first. Timings include writing the output to disk, but we checked the outputs outside the timer before deleting them. We left the page cache in place between runs (`DROP_CACHES=0`), so timings may reflect input cached by earlier runs.

The repeat spread of each combination of dataset, tool, format and thread count is (maximum − minimum) / median of the wall times of its two runs. The median spread over all combinations was 0.8%. At eight threads, no combination with untagged gzip FASTQ had a spread of more than 1.1%, and no combination with tagged gzip FASTQ had a spread of more than 1.4%.

Larger spreads occurred in some plain FASTQ runs because of the page cache. On plain ligation FASTQ at one and four threads, whittle read the file from disk in all four of its runs. chopper found most of the file in the cache in the first repetition and read it from disk in the second. When reading from disk, chopper took 70.10 and 47.61 seconds with one and four threads, respectively, and whittle took 55.25 and 38.46 seconds. When reading most of the file from the cache, chopper took 33.34 seconds with one thread, giving a spread of 71.1%. chopper had the shorter two-run median at both thread counts: 51.7 against 56.6 seconds with one thread and 35.7 against 39.5 seconds with four. whittle used less CPU time in both repetitions: 29 seconds with one thread and 38 to 39 seconds with four, compared with 33 to 39 seconds and 58 to 64 seconds for chopper.

**Figure 1E.** On plain FASTQ (top plot), fastplong's median wall time fell from one to eight threads and rose again at 16 threads, while on gzip FASTQ (bottom plot) it changed little with the thread count. Per-dataset values are in the archived results.

Table S7 gives the eight-thread medians for all five formats. On untagged gzip FASTQ at eight threads, whittle's median memory peaks were at most 142 MiB, and fastplong's largest, on ultra-long reads, was 15,187 MiB. chopper's gzip pipeline had median peaks between 34 MiB (ligation) and 8,151 MiB (SPRQ-Nx). On Fiber-seq, whittle took longer on the uBAM than on the FASTQ files exported from it. The uBAM is a 51.9 GiB file with the four kinetic arrays in every record.

**Table S7.** Two-run medians of wall time, CPU time and peak memory for cropping and filtering at eight threads. The Input and output column gives the format read and written by a run, and the FASTQ and gzip FASTQ rows refer to untagged files. Wall is the wall time and CPU is the user plus system time, both in seconds. Memory is the peak memory in MiB. For chopper with pigz, CPU is summed over both programs and Memory is the larger of their peaks. n/a marks a format that a tool was not run on, because chopper and fastplong cannot read uBAM and fastplong cannot rewrite modification tags. A ratio is n/a if it needs such a run. The whittle ratios columns give the speed ratios and CPU-time percentages reported in the main text. Speed vs chopper and Speed vs fastplong are the wall time of that tool divided by whittle's, so a value above 1 means that whittle was faster. CPU (% of chopper) is whittle's CPU time as a percentage of chopper's. These values are calculated from unrounded medians, so they can differ slightly from ratios of the rounded medians. For the three tools, bold marks the lowest wall time, CPU time and memory in each dataset and format, and ties at the displayed precision are all bold. If only one tool was run on a dataset and format, its values are not shown in bold. In the whittle ratios columns, bold marks the highest value of each format across the four datasets.

| | | whittle | | | chopper | | | fastplong | | | whittle ratios | | |
|---|---|---|---|---|---|---|---|---|---|---|---|---|---|
| **Dataset** | **Input and output** | **Wall (s)** | **CPU (s)** | **Memory (MiB)** | **Wall (s)** | **CPU (s)** | **Memory (MiB)** | **Wall (s)** | **CPU (s)** | **Memory (MiB)** | **Speed vs chopper** | **Speed vs fastplong** | **CPU (% of chopper)** |
| ONT ligation | FASTQ | **45** | **42** | 104 | 49 | 71 | **25** | 73 | 540 | 6,301 | 1.1 | 1.6 | 59.4 |
| | gzip FASTQ | **89** | **538** | 106 | 120 | 988 | **34** | 456 | 1,082 | 6,363 | **1.4** | 5.1 | 54.4 |
| | tagged FASTQ | **87** | **94** | 104 | 103 | 220 | **56** | n/a | n/a | n/a | 1.2 | n/a | 42.7 |
| | tagged gzip FASTQ | **100** | **586** | 107 | 131 | 1,195 | **32** | n/a | n/a | n/a | 1.3 | n/a | 49.0 |
| | uBAM | 102 | 567 | 35 | n/a | n/a | n/a | n/a | n/a | n/a | n/a | n/a | n/a |
| ONT ultra-long | FASTQ | **47** | **38** | 178 | 56 | 57 | **116** | 213 | 1,688 | 14,362 | 1.2 | **4.6** | **66.9** |
| | gzip FASTQ | **85** | **553** | 142 | 113 | 949 | **61** | 470 | 2,009 | 15,187 | 1.3 | 5.5 | 58.3 |
| | tagged FASTQ | **63** | **81** | 195 | **63** | 188 | **107** | n/a | n/a | n/a | 1.0 | n/a | 43.3 |
| | tagged gzip FASTQ | **94** | **610** | 169 | 131 | 1,155 | **55** | n/a | n/a | n/a | **1.4** | n/a | 52.8 |
| | uBAM | 94 | 532 | 78 | n/a | n/a | n/a | n/a | n/a | n/a | n/a | n/a | n/a |
| Revio Fiber-seq | FASTQ | **26** | **22** | 116 | 34 | 37 | **54** | 45 | 376 | 1,359 | **1.3** | 1.8 | 59.1 |
| | gzip FASTQ | **42** | **291** | **130** | 51 | 475 | 4,495 | 242 | 613 | 1,357 | 1.2 | **5.8** | **61.3** |
| | tagged FASTQ | **26** | **43** | 159 | 32 | 77 | **71** | n/a | n/a | n/a | 1.2 | n/a | **55.4** |
| | tagged gzip FASTQ | **43** | **299** | **159** | 53 | 539 | 3,415 | n/a | n/a | n/a | 1.2 | n/a | **55.4** |
| | uBAM | 179 | 918 | 73 | n/a | n/a | n/a | n/a | n/a | n/a | n/a | n/a | n/a |
| Revio SPRQ-Nx | FASTQ | **52** | **41** | 146 | 66 | 66 | **75** | 69 | 525 | 1,011 | **1.3** | 1.3 | 61.6 |
| | gzip FASTQ | **62** | **442** | **132** | 79 | 743 | 8,151 | 357 | 918 | 1,194 | 1.3 | 5.7 | 59.5 |
| | tagged FASTQ | **44** | **100** | 169 | 81 | 296 | **91** | n/a | n/a | n/a | **1.9** | n/a | 33.8 |
| | tagged gzip FASTQ | **68** | **474** | **131** | 84 | 1,051 | 4,257 | n/a | n/a | n/a | 1.2 | n/a | 45.1 |
| | uBAM | 76 | 398 | 25 | n/a | n/a | n/a | n/a | n/a | n/a | n/a | n/a | n/a |

**Output checks.** seqkit summarised the output of every run. Sequali reported on the uBAM, gzip FASTQ and tagged gzip FASTQ versions of each input. For each dataset, format and tool, it also reported on the output of the first repetition at 16 threads. A tool's output on a given dataset and format had the same read and base counts in both repetitions and at all four thread counts. For the gzip FASTQ outputs, Table S8 lists these counts and the N50. The Sequali reports, the per-run statistics and the timing logs are archived in full.

whittle kept 94.83% of the input bases from the ligation reads, 97.83% from the ultra-long reads, 99.91% from the Fiber-seq reads and 99.89% from the SPRQ-Nx reads. chopper's output matched whittle's in read and base counts on the ultra-long, Fiber-seq and SPRQ-Nx inputs. From the ligation input, whittle kept 1,601 more reads and 3,511,557 more bases than chopper, consistent with chopper's filtering before cropping. fastplong kept 72,384 more reads than whittle from the ligation input and 2,342 more from the ultra-long input, which fits its use of the arithmetic mean of Phred scores.

**Table S8.** Read and base counts of the inputs and outputs of gzip FASTQ cropping and filtering, with the N50 of each output. Output counts were identical at every thread count tested and in both repetitions. Base counts are in Gb ($10^9$ bases). The benchmark reports hold the statistics of the other formats, the read qualities and the GC content.

| Dataset | Tool | Input reads | Output reads | Input Gb | Output Gb | Output N50 (bp) |
|---|---|---|---|---|---|---|
| ONT ligation | whittle | 8,030,530 | 4,118,443 | 29.980 | 28.430 | 18,642 |
| ONT ligation | chopper | 8,030,530 | 4,116,842 | 29.980 | 28.427 | 18,643 |
| ONT ligation | fastplong | 8,030,530 | 4,190,827 | 29.980 | 28.869 | 18,631 |
| ONT ultra-long | whittle | 216,065 | 211,883 | 30.001 | 29.350 | 149,052 |
| ONT ultra-long | chopper | 216,065 | 211,883 | 30.001 | 29.350 | 149,052 |
| ONT ultra-long | fastplong | 216,065 | 214,225 | 30.001 | 29.660 | 148,945 |
| Revio Fiber-seq | whittle | 745,544 | 745,540 | 16.941 | 16.926 | 22,764 |
| Revio Fiber-seq | chopper | 745,544 | 745,540 | 16.941 | 16.926 | 22,764 |
| Revio Fiber-seq | fastplong | 745,544 | 745,540 | 16.941 | 16.926 | 22,764 |
| Revio SPRQ-Nx | whittle | 1,676,535 | 1,676,530 | 30.015 | 29.982 | 19,162 |
| Revio SPRQ-Nx | chopper | 1,676,535 | 1,676,530 | 30.015 | 29.982 | 19,162 |
| Revio SPRQ-Nx | fastplong | 1,676,535 | 1,676,530 | 30.015 | 29.982 | 19,162 |

### S5 Tool capabilities

Table S9 compares the tools on uBAM input, annotation updates, chimera splitting and adapter finding. The capability grid of Figure 1B is drawn from this table.

**Table S9.** Capabilities of whittle and related tools, with their sources. Each entry describes the listed version of a tool. For modkit repair, the entries describe its documentation as read on 23 October 2026. The Evidence column gives the sources behind each row. The MM/ML, Moves and Kinetics columns state whether trimming updates the annotation. The entry FASTQ marks updates in tagged FASTQ only, and n/a marks an annotation of the other platform. whittle's move-table updates require `--update-moves` and uBAM output. The Splits chimeras column refers to splitting at adapters, and the Finds adapters column refers to discovery when no sequences are supplied. The last two rows, dorado basecaller 2.1.0 and modkit repair, are not shown in Figure 1B.

| Tool | uBAM input | MM/ ML | Moves | Kinetics | Splits chimeras | Finds adapters | Evidence |
|---|---|---|---|---|---|---|---|
| whittle | yes | yes | yes | yes | yes | yes | Section S2 |
| chopper 0.14.1 | no | FASTQ | no | no | no | no | README, `--update-mods` |
| dorado trim 2.1.2 | yes | yes | yes | n/a | no | no | source, `demux/Trimmer.cpp` |
| lima 26.2.1 | yes | yes | n/a | yes | no | no | changelog 2.5.0, 2.6.0 and 2.12.0, pbbioconda issue 610, tested on synthetic reads; Iso-Seq mode in Section S4.4 |
| fastplong 0.7.1 | no | no | no | no | yes | yes | `--help` and README |
| Porechop_ABI 0.5.1 | no | no | no | no | yes | yes | Bonenfant et al. 2023, README and source |
| cutadapt 5.2 | yes | no | no | no | no | no | release notes, version 4.7 |
| dorado basecaller 2.1.0 | no | yes | yes | n/a | yes | no | read splitting documentation |
| modkit repair | yes | yes | no | n/a | no | no | repair documentation |
